# Interspecies Organoids Reveal Human-Specific Molecular Features of Dopaminergic Neuron Development and Vulnerability

**DOI:** 10.1101/2024.11.14.623592

**Authors:** Sara Nolbrant, Jenelle L. Wallace, Jingwen W. Ding, Tianjia Zhu, Jess L. Sevetson, Janko Kajtez, Isabella A. Baldacci, Emily K. Corrigan, Kaylynn Hoglin, Reed McMullen, Megan S. Ostrowski, Matthew T. Schmitz, Arnar Breevoort, Dani Swope, Fengxia Wu, Bryan J. Pavlovic, Sofie R. Salama, Agnete Kirkeby, Hao Huang, Nathan K. Schaefer, Alex A. Pollen

## Abstract

The disproportionate expansion of telencephalic structures during human evolution involved tradeoffs that imposed greater connectivity and metabolic demands on midbrain dopaminergic neurons. Despite the central role of dopaminergic neurons in human-enriched disorders, molecular specializations associated with human-specific features and vulnerabilities of the dopaminergic system remain unexplored. Here, we establish a phylogeny-in-a-dish approach to examine gene regulatory evolution by differentiating pools of human, chimpanzee, orangutan, and macaque pluripotent stem cells into ventral midbrain organoids capable of forming long-range projections, spontaneous activity, and dopamine release. We identify human-specific gene expression changes related to axonal transport of mitochondria and reactive oxygen species buffering and candidate *cis-* and *trans*-regulatory mechanisms underlying gene expression divergence. Our findings are consistent with a model of evolved neuroprotection in response to tradeoffs related to brain expansion and could contribute to the discovery of therapeutic targets and strategies for treating disorders involving the dopaminergic system.

## Introduction

Midbrain dopaminergic (DA) neurons coordinate multiple aspects of cortical- and subcortical functions and regulate motor control, as well as recently evolved human cognitive and social behaviors^1^. Dysregulation and degeneration of these neurons are major drivers in various disorders that are unique or enriched in humans, including schizophrenia and Parkinson’s disease (PD)^2,3^, but the origins of human-specific developmental trajectories and vulnerabilities remain poorly understood.

Evolutionary tradeoffs may contribute to the increased connectivity requirements and vulnerabilities of human DA neurons^4,5^. The evolutionary expansion of the human brain disproportionately increased the size of DA target regions in cortex and striatum compared to source regions in the ventral midbrain^1,4,6–9^. In total, 400,000 DA neurons in substantia nigra and ventral tegmental area must supply dopamine to expanded human forebrain structures, with each human midbrain DA neuron predicted to form over 2 million synapses^1,10,11^. In addition, the density of DA afferents in basal ganglia has also increased in humans compared to chimpanzees^12,13^, and great ape axons show more complex morphology and altered cortical target distribution^14^. One recent model suggests that the evolution of cooperative behaviors in humans required increased DA innervation in the medial caudate and nucleus accumbens^12,15^. Thus, the human DA system may be under pressure both to adapt to innervating a large forebrain and to regulating evolutionarily novel behaviors.

DA neurons are under intense mitochondrial and bioenergetic demands, which are thought to contribute to their selective vulnerability to degeneration in PD^16,17^. Midbrain DA neurons are unmyelinated, long-projecting, highly arborized cells that release dopamine, the production of which generates toxic reactive oxygen species byproducts^18,19^. Pacemaker activity through L-type calcium channels further increases the energetic demands on DA neurons as a high cytosolic concentration of calcium requires additional ATP for extrusion and can lead to mitochondrial damage^20–22^. Several pathways have been proposed as protective in DA neurons, including cell-intrinsic buffering of free radicals and calcium^23,24^. We hypothesize that the utilization of these or other protective mechanisms could have increased in human neurons compared to other primates, as compensatory adaptations in response to increased DA connectivity requirements.

In this study, we established a phylogeny-in-a-dish approach, generating interspecies ventral midbrain cultures of human, chimpanzee, orangutan and macaque cell lines, and performed combined single nucleus (sn)RNA sequencing and assay for transposase-accessible chromatin (ATAC) sequencing during midbrain progenitor specification in 2D cultures (day 16) and maturation in organoids (day 40-100). Although developing tissue from apes is inaccessible for ethical reasons, these pluripotent stem cell (PSC)-based approaches enable comparative developmental studies and isolation of cell-intrinsic species differences^25,26^. By comparing homologous cell types across primate species, we discover increased expression in the human lineage of genes related to axonal transport of mitochondria and reactive oxygen species buffering, candidate *cis*-regulatory mechanisms enriched in noncoding structural variants, and *trans*-regulatory mechanisms involving gene networks driven by DA lineage-enriched transcription factors (TFs) OTX2, PBX1, and ZFHX3. Subjecting interspecies organoids to rotenone-induced oxidative stress^27,28^ unmasked human-specific responses, consistent with a model of increased neuroprotective mechanisms in human DA neurons. Together, these findings provide a comparative multiomic atlas of primate ventral midbrain differentiation and implicate candidate molecular pathways supporting DA neuron specializations in the enlarged human brain.

## Results

### PSC-derived interspecies cultures model primate ventral midbrain specification and development

We first considered species differences in neuroanatomical scaling influencing the cellular environment of DA neurons. While stereological surveys provide an account of species difference in DA neuron number^8,9^, we further applied a standardized approach to quantify differences in target region volume and axon tract length across multiple adult human and macaque individuals. Magnetic resonance imaging (MRI) revealed an 18-fold expansion of the prefrontal cortex (PFC) and a 6.8-fold expansion of striatum target region volumes in humans (Figures 1A, 1B and Table S1). Similarly, diffusion tensor imaging (DTI), indicated a 2.5 fold increase in fiber tract length from DA nucleus substantia nigra to the caudate (Figures S1A and S1B). Although DTI cannot quantify differences in axonal arborization that represents the majority of axon length, these results further support the increased connectivity requirements of midbrain DA neurons throughout the extended human lifespan (Figure 1C) in the expanded human brain. The DA innervation densities in these target structures have been carefully investigated in previous studies ^12,13^, that have shown a human-specific increase in DA innervation density in parts of the basal ganglia which should further exacerbate the burden on human DA neurons.

**Figure 1:**
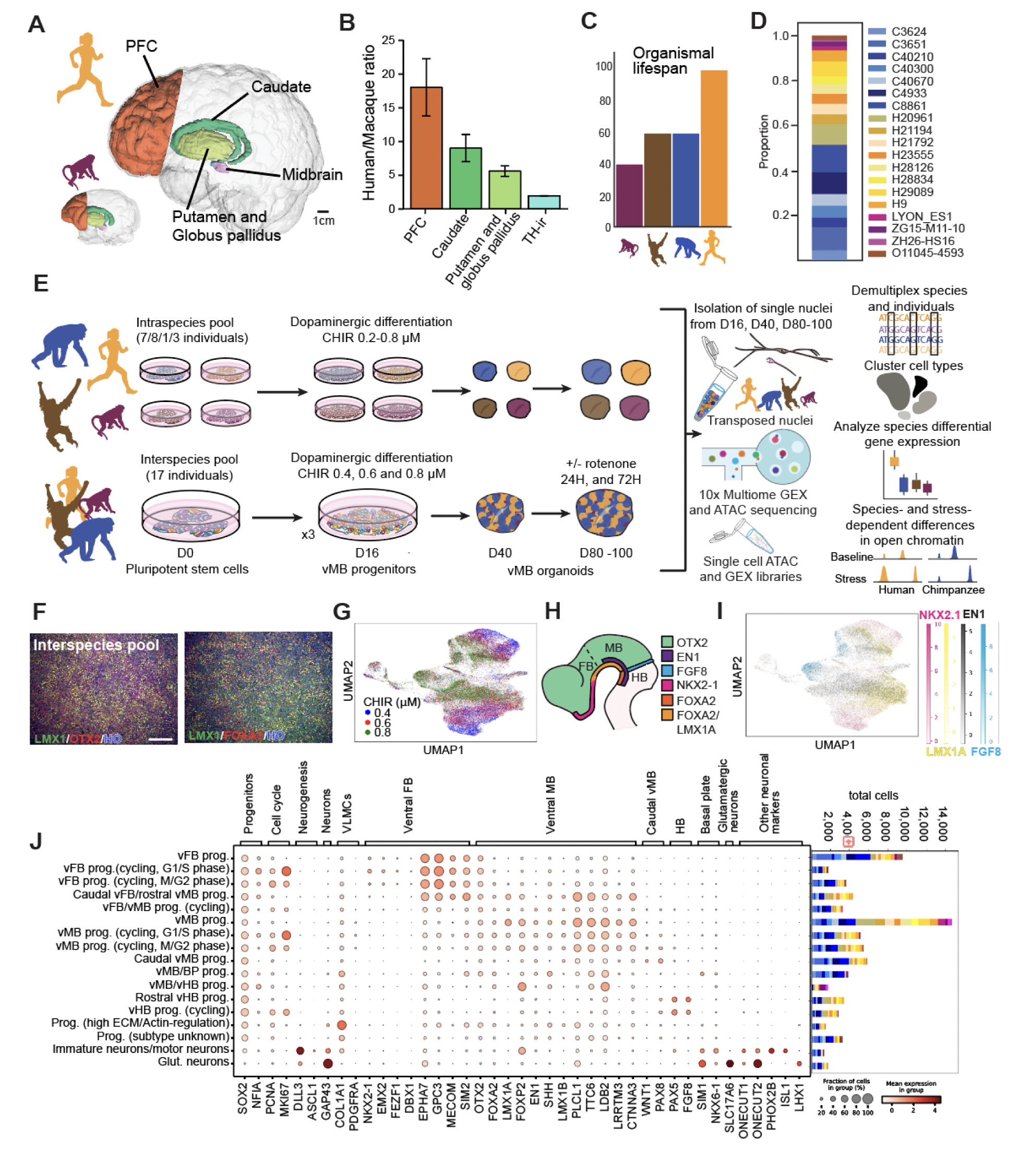
Pluripotent stem cell-derived interspecies cultures model primate ventral midbrain specification and development. A. Unequal scaling of dopamine target regions in human compared to macaque with regions quantified by MRI highlighted. B. Human/macaque volume ratios from MRI quantification of dopamine target regions PFC (19.52 ± 4.24, p=3.33E-08), caudate (9.17 ± 2.01, p=5.56E-11), and putamen/globus pallidus (5.68 ± 0.78, p=3.54E-09, n=10 human individuals, n=3 macaque individuals). The ratio of DA neuron numbers was calculated through comparison of TH-ir cells stereotactically quantified by^8,9^ (1.93 ± 0.024, p=4.95E-07, n=6 human individuals, n=7 macaque individuals). C. An increased organismal lifespan^101,102,103^ is an additional factor that contributes to the enhanced stress human DA neurons face. D. Proportions of all individuals at D16, post quality control and doublet removal. Human lines: H9, H20961, H21194, H21792, H23555, H28126, H28334, H29089; Chimpanzee lines: C3624, C3651, C4933, C8861, C40210, C40300, C40670; Orangutan line: O11045-4593; Macaque lines: LYON-ES1, ZG15-M11-10, ZH26-HS16. E. Experimental design for generating inter- and intraspecies midbrain organoids for paired snRNA- and ATAC-sequencing. F. Interspecies ventral midbrain culture at D14, with immunocytochemical labeling of FOXA2, OTX2 and LMX1A/B. s. G. UMAP of cells collected at D16, colored by MULTI-seq barcode identity which corresponds to the applied CHIR99021 concentration. H. Rostral-caudal expression domains of key genes to determine cell identity within ventral diencephalon, midbrain and hindbrain. I. UMAP of cells collected at D16, colored by key genes to determine rostral-caudal identity. J. Dotplot of the expression of cell type markers for the assigned cluster identities at D16 (left), with a bar chart of the contribution of individuals to the different cell types (right), with colors as labeled in D. Scale bars: 200 μm. For the bar chart in Figure 1A, data are represented as mean ± standard deviation. See also Figure S1 and S2. PFC, prefrontal cortex; TH, Tyrosine hydroxylase; prog, progenitors; vMB, ventral midbrain; vFB, ventral forebrain, vHB, ventral hindbrain; BP, basal plate; ECM, extracellular matrix; glut, glutamatergic; DA, dopaminergic; VLMCs, vascular leptomeningeal cells.

We next established cellular models to explore developmental gene regulatory programs that may contribute to species differences in connectivity and cellular vulnerability. To study divergence in gene expression and chromatin accessibility during primate midbrain specification and differentiation, we developed a phylogeny-in-a dish approach where multiple PSC-lines (human=8, chimpanzee=7, orangutan=1, and macaque=3, Figure 1D, see Table S2 for experimental details) were differentiated together into ventral midbrain progenitors as either intra- or interspecies pools^27,29,30^, using an established protocol^31^ that produces functional DA neurons, validated in preclinical models of PD^32–34^, adapted to 3D maturation conditions (Figures 1E and S1C). Pooling many PSC-lines for differentiation provides an efficient strategy to isolate cell-intrinsic differences that can be attributed to species, rather than cell lines, batch effects, or culture conditions.

In vitro specification to ventral midbrain identities recapitulates the developmental morphogen environment and relies on Sonic hedgehog (SHH) signaling for ventralization to a floor plate identity and activation of WNT signaling, by the addition of the GSK3 inhibitor CHIR99021 (CHIR), to obtain midbrain fates. To generate cell type identities corresponding to the rostro-caudal extent of the developing midbrain, including the caudal diencephalon and rostral hindbrain^35^, we employed three different concentrations of CHIR in parallel pools and used immunocytochemistry to confirm patterning to ventral midbrain identities (Figures 1F and S1D-S1F, and S1G-J for additional outgroup individuals).

After verifying overall patterning, we proceeded to measure gene expression and chromatin accessibility in single cells via snRNA-and snATAC-sequencing (10x Genomics Multiome kit) from the different pools (Figure S2A and Table S2). Following the removal of doublets and low quality cells, 73,077 nuclei were retained for downstream analysis across three experiments, 38,066 from interspecies pools and 35,011 from intraspecies pools (Figure S2B and Table S2). Demultiplexing species and individual identity revealed comparable numbers of human (30,543) and chimpanzee (37,523) cells, and preservation of all individuals in intra- and interspecies pools (Figures 1D and S2E-S2G).

Principal component analysis (PCA) of the gene expression data at day 16 (D16) revealed species and maturation/cell cycle stage as the main drivers of separation along PC1 and PC2, respectively (Figures S2H and S2I). To identify homologous cell types between species, batch integration of batch-balanced k-nearest neighbors (BBKNN) was applied, resulting in Leiden clusters with mixed species contributions (Figures S2C, S2D and S2J). Labeling cells based on pool-of-origin CHIR concentration using MULTI-seq barcodes^36^ revealed a gradient of CHIR concentrations across the UMAP (Figures 1G), which corresponded to a gradient of genes with defined rostral to caudal expression patterns (Figures 1H and 1I). Consensus non-negative matrix factorization (cNMF)^37^ indicated that increasing CHIR concentration shifted cells from rostral to caudal ventral midbrain progenitor identities across species (Figures S2O-S2Q).

Using cluster-enriched genes and known markers from the literature, we assigned cell type identities to each cluster and confirmed the presence of progenitor cell types ranging from diencephalon to hindbrain, with ventral midbrain progenitors being the most abundant (Figures 1J and S2K). At this stage of differentiation, the majority of cells were still proliferative progenitors, with a smaller subset of postmitotic neurons (Figure S2L). The expression of regional marker genes supported the equivalent response of human and chimpanzee lines to the patterning protocol (Figures S2M and S2N). Additionally, all individuals from all four species (with the exception of one macaque line) were represented in each cell type cluster (Figure 1J and Table S2).

In summary, phylogeny-in-a-dish culture enables equivalent patterning of human and chimpanzee PSCs to homologous progenitor subtypes, including ventral midbrain progenitors, and allows for comparative analysis of molecular programs underlying early stages of specification.

### Transcriptional landscape, reproducibility, and fidelity of interspecies organoids

To study the development and maturation of DA neurons and related neuronal subtypes, D16 progenitors were aggregated into neurospheres for continued culture in a 3D environment (Figures S1K, S1L, S1N and S1O). Immunohistochemistry revealed abundant DA neurons in all four species (Figures 2A, S1N, S1P, S1Q and S1S) and in interspecies pools (Figures 2B, S1M, S1O and S1R), with a predominance of TH^+^ axons at the periphery. DA neurons in D40 organoids formed TH^+^ projections that innervated fused cortical organoids in mesocortical assembloids (Figure S1T). High-density multi-electrode array (HD-MEA) recordings initially showed mainly uncoordinated, tonic spiking at D60, with some coordinated burst activity emerging in both human and chimpanzee organoids cultured on the HD-MEA around D70 (Figures S3A and S3C). By D90, bursts (clear groupings separated by less active periods, characterized by bimodal inter-burst intervals) emerged in human and chimpanzee organoids (Figures S3B and S3D). At this point, bursts recruited most detected neural units, and tended to last 0.5-1.5 seconds (Figures S3B, S3D and S3E-F). Spontaneous dopamine release could be detected by D30, increased over the next few weeks, and then remained stable (Figures S3G-I). These analyses suggest that ventral midbrain organoids recapitulate important structural and functional aspects of normal DA neuron development.

**Figure 2:**
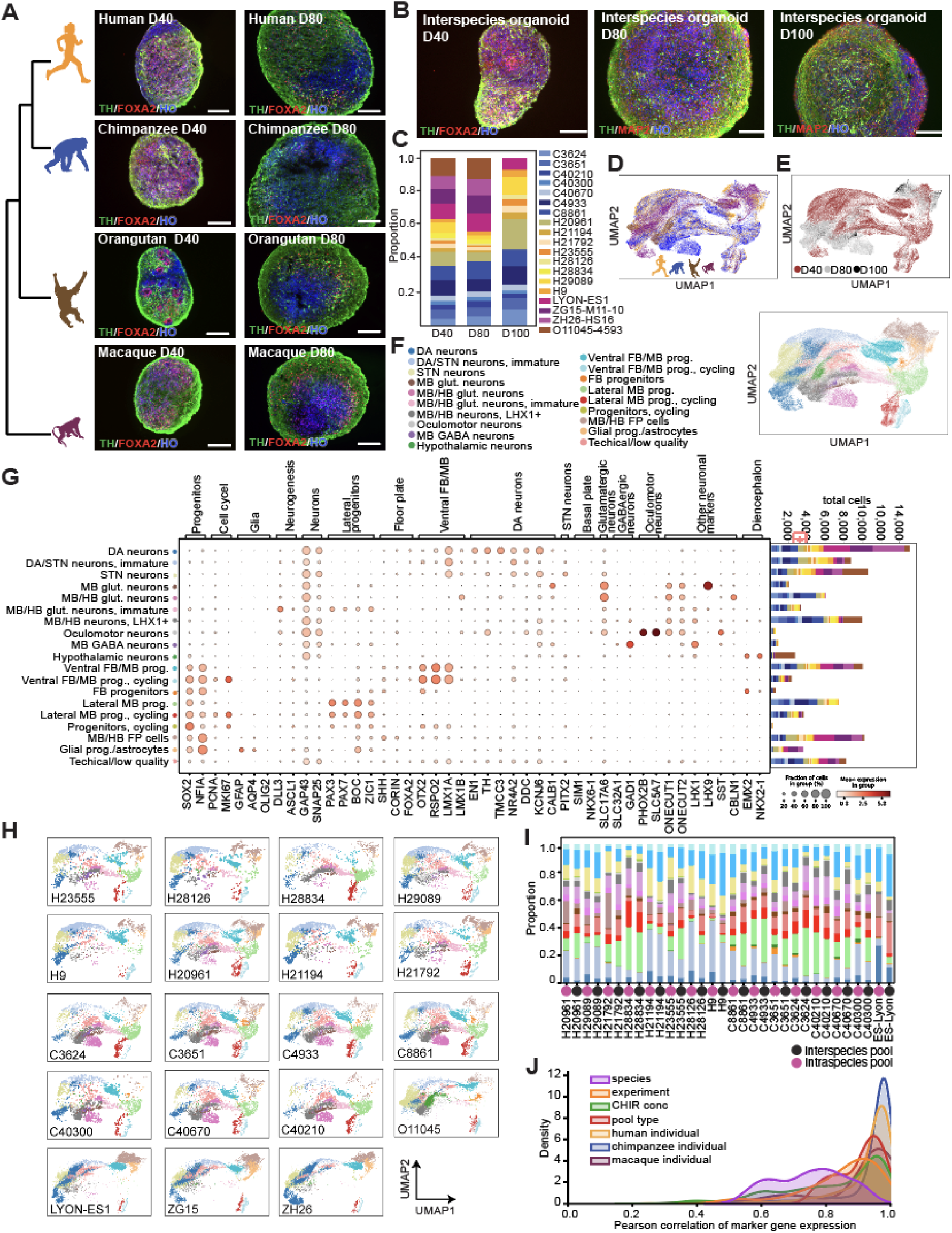
Transcriptional landscape, cell type diversity and reproducibility of developing primate ventral midbrain organoids. A. Ventral midbrain organoids from the human pool (n=8 individuals), chimpanzee pool (n=7 individuals), orangutan (O11045-4593), and macaque (LYON-ES1), labeled for TH, FOXA2 and Hoechst at D40 and D80. B. Interspecies ventral midbrain organoids, labeled for TH/FOXA2 or TH/MAP2 and Hoechst at D40, 80 and 100. C. Proportions of all individuals at D40, 80 and 100, post quality control and doublet removal. D-F. UMAPs of cells collected at D40-100, colored by species (D), timepoint (E) and assigned cell type identity (F). G. Dotplot of the expression of cell type markers for the assigned cluster identities at D40-100 (left), with a bar chart of the contribution of individuals to the different cell types (right), with colors as labeled in C. H. UMAPs for each individual, colored by cell type identity. I. Cell type proportions at D40, for each individual where data from both interspecies and intraspecies pools were collected. J. Histograms summarize distribution of cell type correlations across marker genes between conditions for each level of experimental design and replication for D40-100 cell types. Cell type transcriptomes are highly correlated for interindividual comparisons within species, comparisons across CHIR conditions, and comparisons across pool types. Note that the increased divergence across species likely represents real biological variation (as observed *in vivo*^41^) and is greater than that from all other levels of experimental design, consistent with the overall reproducibility of our study and our power to discover species differences in gene regulation. Scale bars: 200 μm. See also Figure S1, S2 and S3. prog, progenitors; vMB, ventral midbrain; vFB, ventral forebrain, vHB, ventral hindbrain; BP, basal plate; ECM, extracellular matrix; glut, glutamatergic; GABA, GABAergic; DA, dopaminergic; STN, subthalamic nucleus; FP, floor plate.

While the multiome data from D16 progenitors captured early stages of cell type specification and commitment, measuring gene expression and chromatin accessibility from organoid stages (D40-100) enables comparative studies of the molecular basis of later developmental events including axonogenesis, neurotransmitter synthesis, and the onset of neuronal activity. To increase the temporal resolution of the maturation process, data was collected from D40, 80 and 100 (Figure 2E). After removing low quality cells, doublets, and clusters defined by high endoplasmic reticulum (ER) stress^38,39^, 105,777 cells remained for clustering and downstream analysis. Even after extended organoid culture, all human and chimpanzee individuals were recovered from both intra- and interspecies pools (Figures 2C, 2D and S2E-G; total number of cells: chimpanzee=39,300, human=27,293). Due to loss of cells from outgroup species in the interspecies pool, additional samples of macaque and orangutan (contributing to a total of macaque=27,969 and orangutan=11,215 cells), maintained and matured in parallel with thawed replicate samples of human and chimpanzee intraspecies pools were added from subsequent differentiations (Table S2).

BBKNN integration and Leiden clustering resulted in 30 mixed species clusters (Figures S2R and S2S). Cell type identities were manually assigned using cluster-enriched genes and known markers (Figures 2F, 2G and S2T-X), and constituted a mix of both species and individual identities (Figure 2G). While human and chimpanzee progenitors gave rise to a range of neuronal subtypes including DA, glutamatergic and GABAergic neurons (Figures S2U and S2V), the macaque lines predominantly yielded cells of the DA lineage, and contributed less to the other cell type clusters (Figures 2G and S2X). Interestingly, there was a cluster of immature neurons with mixed DA/STN marker expression which split into separate DA and STN neuron clusters with maturation, consistent with previously described transcriptional similarity of these two lineages during development^40^.

We next investigated the reproducibility in cell type compositions across species, individual and pool type. At both D16 (Figure S4A) and D40-100 (Figure 2H), comparable representation of cell types across individuals was observed, highlighting the robustness of the patterning protocol. This reproducibility also extended to pool type, where similar representations of cell type identities between human and chimpanzee individuals were seen at D16 and D40 regardless if the individual was part of an intra- or interspecies pool (Figure 2I, Figure S4B), further supporting the use of our novel interspecies culture system. To quantify reproducibility, we calculated Pearson correlations for cell type marker genes between conditions at each level of experimental design and replication for D40-100 cell types (Figure 2J). Cell type transcriptomes were highly correlated for interindividual comparisons within species (human: r=0.964; chimpanzee: r=0.976; macaque: r=0.961), across CHIR conditions (r=0.880), experiment (r=0.884) and pool types (r=0.917). Importantly, transcriptional divergence across species (r=0.760) was greater than that from all other levels of experimental design, consistent with cross-species biological variation observed in vivo^41^ and supporting the power of our study to discover species differences in gene regulation. Finally, we observed conserved expression of canonical marker genes within the DA lineage at both D16 and D40-100, between species, individuals and pool types, further emphasizing the reproducibility of DA neuron specification across experimental conditions (Figure S4C).

Having established the robustness and reproducibility of our experimental model system, we sought to evaluate the fidelity and maturation of organoid-derived DA neurons. Co-embedding with ventral midbrain data from two primary human datasets^42,43^ revealed co-clustering of homologous cell types (Figures 3A-E, S5A) and shared marker gene expression between primary cells and organoid cells across species (Figure 3F). We next compared the differentiation stage of organoid cells with the absolute age of available primary DA neurons (Figures 3G and 3H). Analysis of shared nearest neighbors among co-embedded cells revealed that maturation in organoids correlates with the age of primary neurons (D40 mean = PCD 51.32, D80 mean = 56.43, D100 mean = 62.72). We further performed pseudotime analysis to compare DA lineage differentiation trajectories between species (Figures 3I-O). These trajectories were similar across pool types and individuals (Figures 3J-L) and reflected the gradient of differentiation stages of captured cells (Figures 3M-O). We observed induction of genes associated with DA neuron maturation, including *KCNJ6* (GIRK2), *TH*, and *SLC8A2.* Interestingly, pseudotime analysis revealed delayed maturation in apes compared with rhesus macaque (Figure 3O), consistent with known differences in gestation time, and a trend for a subtle delay in maturation at D80 in human compared with chimpanzee, consistent with differences observed among cortical neurons^44^. Glial competence, an orthogonal measurement of organoid development, further supported the accelerated differentiation rate in macaque midbrain compared to apes (Figure 3P).

**Figure 3:**
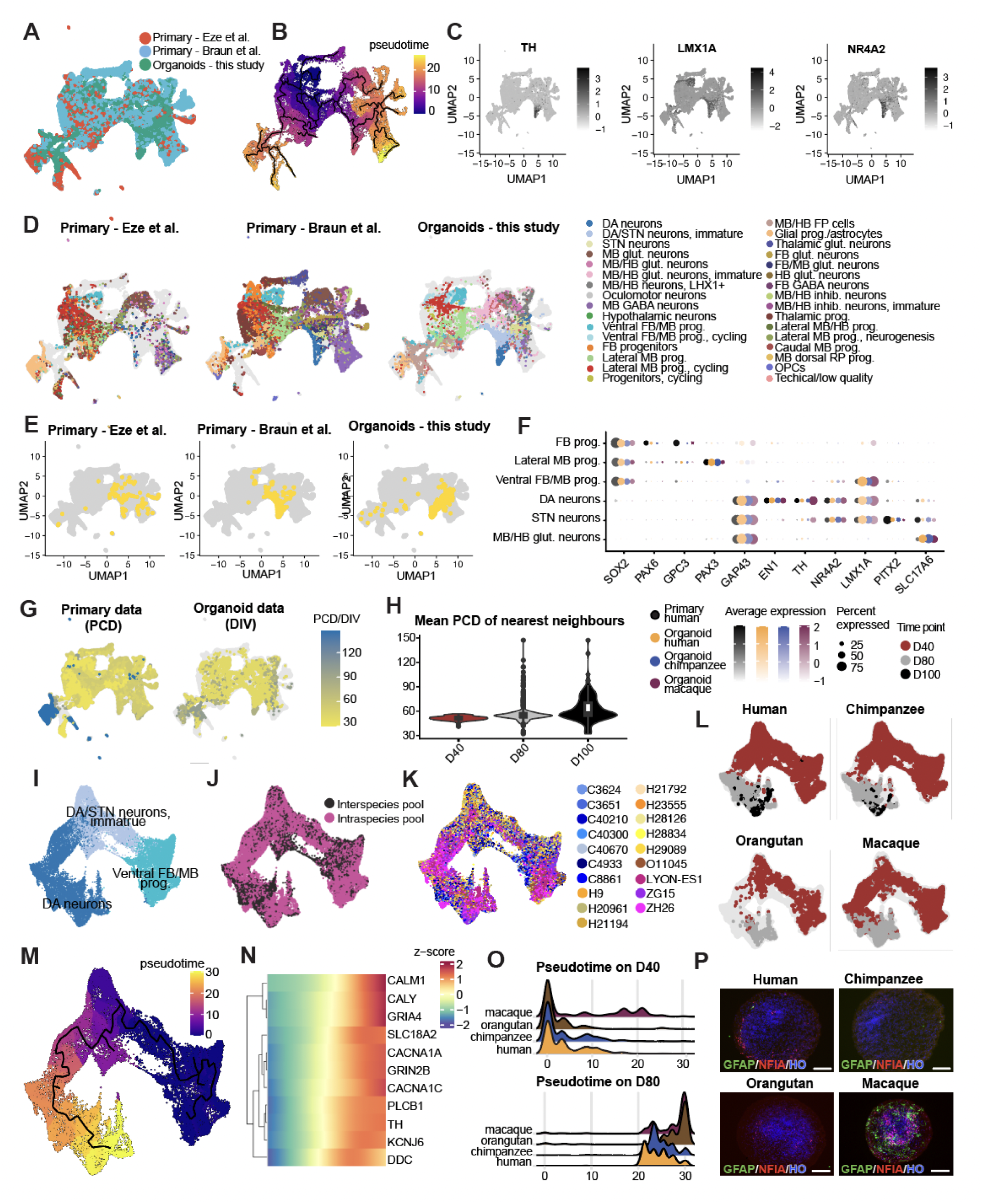
Midbrain organoid fidelity to primary midbrain data and DA neuron maturation timing in human and non-human primates. A. Combined UMAP including two published developing human midbrain datasets from Eze et al (including unpublished data from one additional human individual that we collected) and Braun et al and human midbrain organoids from this study, colored by data source. B. UMAP colored by pseudo time. C. UMAPs colored by gene expression of *TH*, *LMX1A* and *NR4A2*. D. UMAPs colored by supervised cell types, split by data source. E. UMAPs with DA neurons highlighted, split by data source. F. Dotplot for marker gene expression in 6 cell types shared in in vivo primary and in vitro organoid datasets. G. UMAPs colored by timepoint (PCD for primary and DIV for organoids), split by data type H. Violin plot for distribution of mean PCD of nearest primary neighbors of DA neurons in organoids, grouped by timepoint. I-K. UMAP for integrated in vitro organoid DA lineage from 4 species, colored by assigned cell types (I), pool type (J) and individual (K). L. UMAPs of DA lineage cells colored by timepoint and split by species. M. UMAP colored by pseudotime defined within the DA lineage. N. Heatmap for expression of neuronal genes along the DA lineage pseudotime. O. Ridgeplot for pseudotime distribution in DA neurons in 4 species on D40 (top) and D80 (bottom). P. Immunohistochemistry of human pool, chimpanzee pool, macaque (ZH26), and orangutan (O11045) D80 organoids, to visualize GFAP^+^ and NFIA^+^ glial progenitors/astrocytes, counterstained with hoechst (HO). Scale bars: 200 μm. PCD, post conception day; DIV, day in vitro; prog, progenitors; vMB, ventral midbrain; vFB, ventral forebrain, vHB, ventral hindbrain; BP, basal plate; ECM, extracellular matrix; glut, glutamatergic; GABA, GABAergic; DA, dopaminergic; STN, subthalamic nucleus; inhib, inhibitory; FP, floor plate.

In summary, ventral midbrain organoids display reproducible composition and high marker gene conservation across experiments, pool types, individuals, and species. DA neurons show high fidelity to human fetal development, corresponding to midgestation DA neurons, with maturation increasing over time and D100 DA neurons matching the oldest primary human DA neurons collected. Cross-species comparisons highlight comparable cell type diversities and maturation dynamics between human and chimpanzee, enabling the study of species-specific transcriptional differences during the development and maturation of DA neurons and other midbrain cell types.

### Cell type specificity and evolutionary divergence of gene expression across ventral midbrain development

To compare gene expression across species while minimizing the impact of differences in annotation quality, we mapped the snRNA-seq data from all species to an inferred *Homo/Pan* common ancestor genome^45^, with annotations further optimized for single cell transcriptomics analyses^46^. To explore the major sources of gene expression variation in our datasets, we first performed variance partition analysis^47^ on pseudobulk samples for each cell type and gene. Species of origin explained the highest percent of gene expression variance (mean across cell types and genes that met filtering criteria, D16: 19.9%, D40-100: 17.5%), with smaller contributions from individual (cell line) (D16: 7.6%, D40-100: 5.5%), experiment (D16: 4.5%, D40-100: 5.3%), differentiation day (D40-100: 4.9%), pool type (inter-vs intra-species)(D16: 6.4%, D40-100: 1.3%), sequencing lane (D16:1.9%, D40-100: 1.6%) and biological sex (D16: 0.47%, D40-100: 0.36%)(Figure S6A-D).

To compare human and chimpanzee ventral midbrain cell types, we analyzed differentially expressed genes (DEGs) using linear mixed models based on pseudobulk samples across cell types, implemented with the dreamlet package^48,49^ (Figure S6A and S6E-G). We modeled the D16 and D40-100 datasets separately and included the top terms from our variance partition model. As expected, more human-chimpanzee DEGs were detected among more abundant cell types (Figure 4A, dotplot). After filtering for cell types that had at least 200 human and 200 chimpanzee cells, we plotted the correlation of differential expression scores (Methods) for all differentially expressed genes across all cell types (Figure 4A, heatmap). D16 cell types clustered together as did D40-100 cell types, with higher correlations indicating more similar magnitude and direction of differential expression between more closely related cell types.

**Figure 4:**
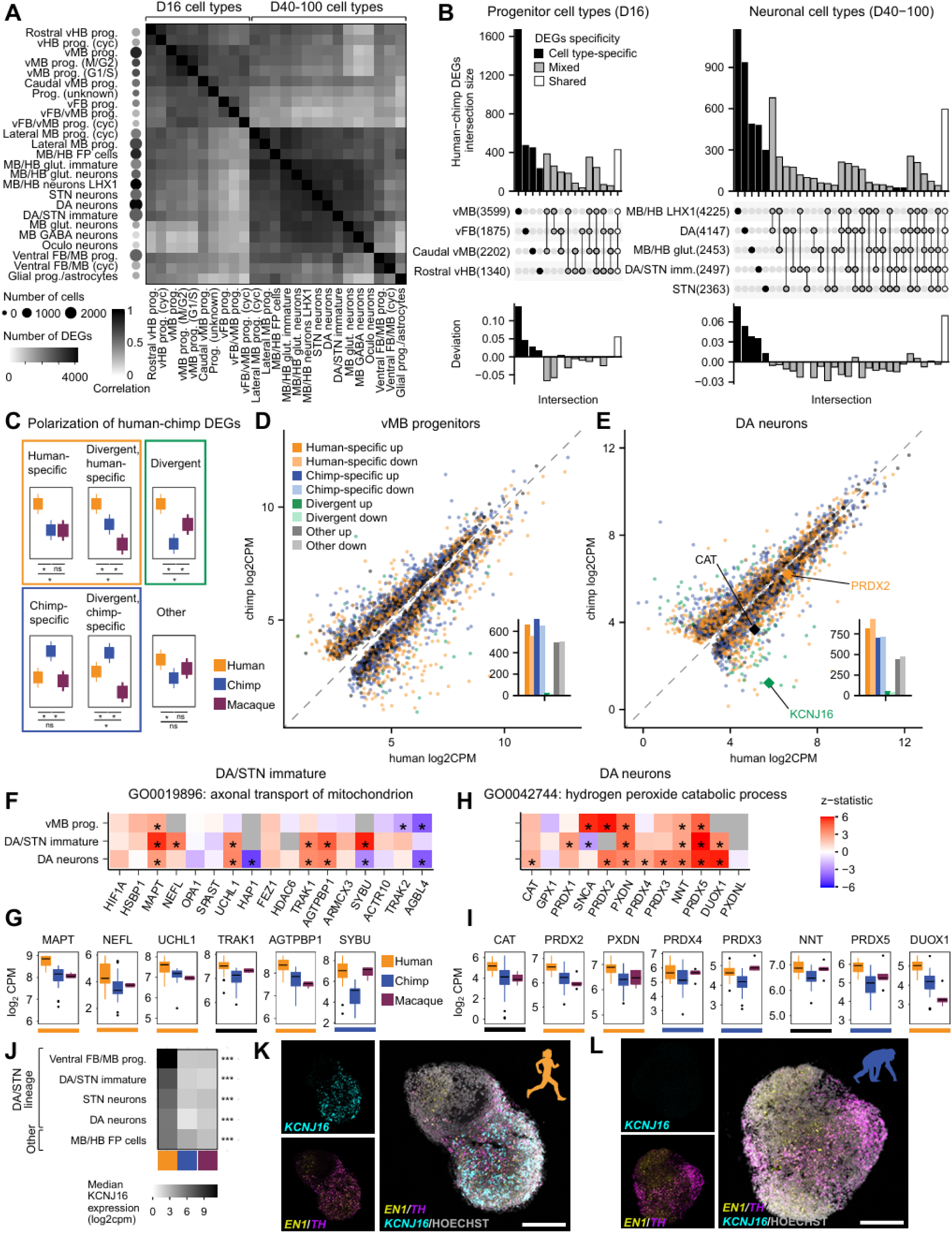
Cell type specificity and evolutionary divergence of gene expression across ventral midbrain development. A. Clustered heatmap showing Pearson correlation between human-chimpanzee DEG scores (logFC * -log10pval) for all genes that were DE in at least one cell type across all D16 and D40-100 cell types with at least 200 cells for both human and chimpanzee. Dotplot on the left shows the number of human and chimpanzee cells (least common denominator) for that cell type, shaded by the number of human-chimpanzee DEGs with FDR < 0.05. B. UpSet plots showing the intersection of human-chimpanzee DEG lists for selected D16 (left) and D40-100 (right) cell types, ordered from cell type specific to shared intersections. Numbers in parentheses represent the total number of human-chimpanzee DEGs for that cell type. C. Scheme for classifying human-chimpanzee DEGs showing which comparisons are significant for each category (*, FDR < 0.05). D. Scatterplot showing average normalized expression across pseudobulk samples for each human-chimpanzee DEG in human versus chimpanzee D16 vMB progenitors, with points colored by categories from C and dotted y = x line. Barplots (insets) show the number of up- and down-regulated genes in each category. E. Same as D for D40-100 DA neurons. F. Heatmap showing z statistics for expressed genes in the top human-upregulated GO term in immature DA/STN neurons. G. Boxplots for human-chimpanzee DEGs belonging to the top human-upregulated GO term in immature DA/STN neurons showing normalized median gene expression values across pseudobulk samples (combination of species, experiment, individual) across species with colored lines below indicating polarization category. H-I. Same as F-G for the top human-upregulated GO term in DA neurons. J. Heatmap showing normalized median expression of *KCNJ16* across human, chimpanzee, and macaque pseudobulk samples. *KCNJ16* expression did not meet the expression threshold in the remaining D40-100 cell types. K-L. RNAscope of *TH*, *EN1* and *KCNJ16* in D40 intraspecies pooled organoids from human (K) and chimpanzee (L). Scale bars: 200 μm DEGs, differentially expressed genes; prog, progenitors; vMB, ventral midbrain; vFB, ventral forebrain, vHB, ventral hindbrain; glut, glutamatergic; GABA, GABAergic; DA, dopaminergic; STN, subthalamic nucleus; Oculo, oculomotor * p < 0.05, ** p < 0.01, *** p < 0.001

To examine the cell type specificity of differential gene expression, we constructed UpSet plots showing the intersection of human-chimpanzee DEGs sets for a subset of progenitor cell types from D16 and neuronal cell types from D40-100. Plotting the deviation for each set intersection showed that there were more cell type-specific and shared human-chimpanzee DEGs than expected based on the number of significant DEGs for each cell type, while DEGs with mixed specificity (shared between some but not all of the plotted cell types) were relatively depleted (Figure 4B).

Next, we focused on the DA lineage (including ventral midbrain (vMB) progenitors (D16), immature DA/STN neurons (D40-100), and DA neurons (D40-100). We examined the evolutionary history of human-chimpanzee DEGs and classified them into four categories based on the significance of gene expression differences in two-way comparisons between human, chimpanzee, and macaque: human-specific (likely derived in the human lineage - human significantly different than chimpanzee and macaque), chimpanzee-specific (chimpanzee significantly different than human and macaque), divergent (all three comparisons significant with macaque gene expression in between human and chimpanzee), and other (human versus chimpanzee is significant but neither species is significantly different from macaque) (Figure 4C, Table S4).These polarization categories could be superimposed on scatterplots showing the average expression of each human-chimpanzee DEG across human versus chimpanzee (Figures 4D, 4E, and S6L) or macaque (Figures S6J, S6K, and S6M) pseudobulk samples in each cell type. In both vMB progenitors and DA neurons, there were similar numbers of human-specific up, human-specific down, chimpanzee-specific up, and chimpanzee-specific down genes (Figures 4D and 4E, insets), supporting our analysis strategy as unbiased. To examine the extent to which organoid models recapitulate species differences, we combined publicly available human and rhesus datasets with newly generated data from 7 individuals (Figures S5A-S5E). The magnitude and direction of differential gene expression in the DA lineage was significantly correlated between human and macaque in vitro models and primary cells across differentiation (Figure S5F) and across species (Figure S5G).

To identify the types of genes that were differentially expressed, we performed competitive gene set analysis on linear mixed model results for human versus chimpanzee in DA lineage cell types. None of the gene sets were significantly enriched at the study-wide FDR (Table S4), reflecting the overall similarity of gene expression in human versus chimpanzee cell types and the low-magnitude, predominantly quantitative expression differences between closely related species. However, several of the top-ranked, nominally significant terms were related to our initial hypotheses about increased connectivity and metabolic demands on human DA neurons. The top two human-upregulated terms in immature DA/STN neurons (but not vMB progenitors or DA neurons) were related to mitochondrion transport: “axonal transport of mitochondrion” (p = 0.003, FDR n.s.) and “mitochondrion transport along microtubule” (p = 0.003, FDR n.s.)(Figure 4F). Moreover, two thirds of the significant human-upregulated genes in this category were classified as human-specific (Figures 4G, S6H, and S6N). In DA neurons, the top human up-regulated term was related to antioxidant activity (“hydrogen peroxide catabolic process”, p = 0.002, FDR = n.s.)(Figure 4H) and several of the significantly human-upregulated genes were human-specific (Figures 4I, S6I, and S6O). Four members of the Peroxiredoxin gene family were expressed at higher levels in human than chimpanzee DA neurons, including human-specific upregulation of *PRDX2*. Interestingly, one of the human-upregulated genes in this set was *CAT* (human vs chimpanzee adj.p.val = 0.039), which encodes the enzyme catalase that degrades hydrogen peroxide and was previously reported to be significantly upregulated in human versus chimpanzee and macaque primary cortical neurons^50^.

Next, we focused on DEGs with high cell type specificity within DA/STN lineage cell types, reasoning that they may have functions specific to the DA system. We calculated the cell type specificity of gene expression across all D40-100 cell types and ranked human-specific and divergent DEGs by their DA lineage specificity scores (Table S4). The top-ranked gene (for both immature DA/STN neurons and DA neurons) was *KCNJ16*, which encodes a pH-sensitive inward-rectifying potassium channel subunit. This gene had highest expression in progenitors but maintained significant expression levels throughout neuronal differentiation, with human expression levels significantly higher than chimpanzee and macaque (Immature DA/STN neurons: human-specific, human vs chimpanzee: logFC = 5.49, p = 6.12e-16, human vs macaque: logFC = 6.92, p = 3.07e-08; DA neurons: divergent, human vs chimpanzee: logFC = 5.06, p = 2.10e-12, human vs macaque: logFC: 3.42, p = 1.64e-06)(Figure 4J). Using RNAscope, we validated human-upregulated *KCNJ16* expression colocalizing with DA markers EN1 and TH (Figures 4K, 4L and S6P-S6R), highlighting a molecular change that could contribute to evolved physiological differences in the DA lineage.

Together, these results represent a resource of human-specific DEGs in ventral midbrain cell types and suggest molecular pathways that may be involved in evolutionary adaptations of the human DA system.

### Evolution of the *cis*-regulatory landscape in ventral midbrain neurons

To explore *cis*-regulatory evolution underlying gene expression differences in ventral midbrain cell types, we turned to the paired snATAC-seq data. To compare open chromatin regions across species, we developed a cross-species consensus peak pipeline, CrossPeak, that focuses on precisely localizing orthologous peak summits across species without introducing species bias. CrossPeak takes as input a set of fixed width, summit-centered peaks for each species and produces a consensus set of peaks across species in the coordinates of each species’ genome as well as retaining species-specific peaks that failed to lift over (Figure 5A, Methods).

**Figure 5:**
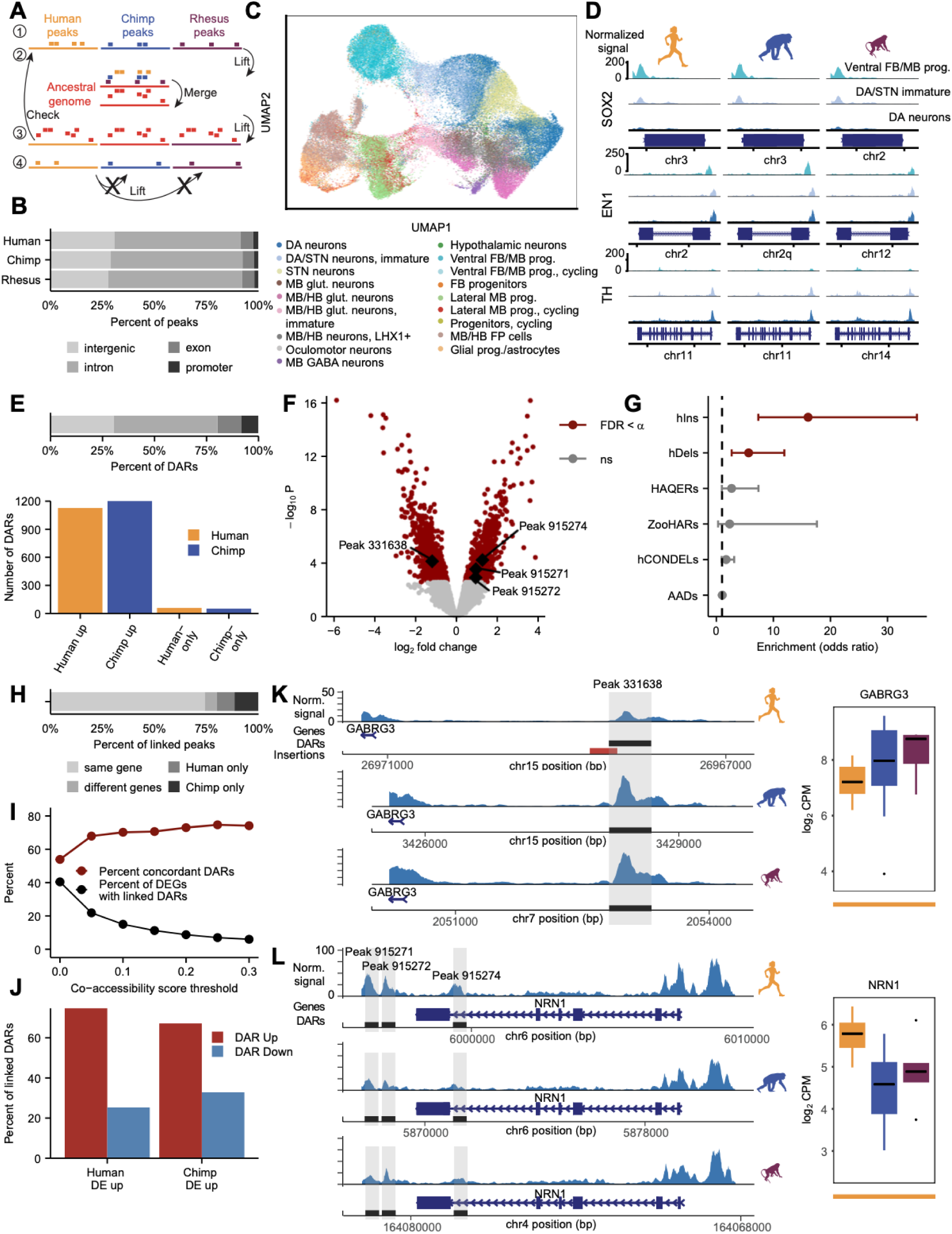
Evolution of the *cis*-regulatory landscape in ventral midbrain neurons. A. Schematic of computational pipeline for identifying consensus and species-specific ATAC-seq peaks across species. Step 1: Use iterative overlap merging to obtain a peak set for each species across all cell types. Step 2: Lift each species peak set to an ancestral genome and merge overlapping peaks according to user-defined rules. Step 3: Lift the consensus peak set back to individual species genomes and check the new position against the original location. Step 4: For any peaks that failed to lift over in steps 2 or 3, attempt to lift them over directly to the other species’ genomes and classify them as species-specific peaks if direct liftover fails. B. Stacked barplot showing the percent of step 1 peaks called within each species located in each genomic category. C. The consensus peak set (step 3 peaks) allows integration of human, chimpanzee, and macaque multiome snATAC-seq data. Cells are colored based on their snRNA-seq cell type annotation. D. Accessibility at marker genes within DA lineage cell types across species. E. Top, stacked barplot showing the percent of human-chimpanzee DARs and species-only peaks in DA neurons located in each genomic category. Bottom, barplot showing the numbers of human-up and chimpanzee-up consensus DARs and the numbers of human-only and chimpanzee-only peaks. F. Volcano plot for human-chimpanzee consensus DARs in DA neurons with alpha = 0.1. G. Forest plot showing odds ratios and confidence intervals for the enrichment of evolutionary signatures within human-chimpanzee DA neuron DARs and species-only peaks with alpha = 0.05. H. Stacked barplot showing the percent of peaks that were linked via Cicero (within DA lineage cell types ventral FB/MB progenitors, DA/STN immature neurons, and DA neurons) to the same gene or different genes within human and chimpanzee data, or were linked only in human or only in chimpanzee. I. Plot showing the relationship between co-accessibility score threshold applied to Cicero links and the percent of DA neuron DEGs with linked DA neuron DARs as well as the percent concordant DARs (defined as the percent of DARs with increased accessibility linked to upregulated DEGs, considering links found in the species where the gene was upregulated). J. Barplot plot showing concordance as the percent of DA neuron DARs linked to upregulated DA neuron DEGs in each species at co-accessibility threshold 0.15. (n = 181 human-up DEGs with 229 linked DARs, 173 of chimpanzee-up DEGs with 231 linked DARs). K. Example of human-downregulated DA neuron DAR linked to human-specific downregulated DA neuron DEG *GABRG3*. Left, pseudobulk DA neuron snATAC-seq signal across human, chimpanzee, and macaque. The DAR overlaps a human-specific insertion from ^51^. Right, boxplot showing normalized *GABRG3* gene expression values for pseudobulk samples across species. Line at bottom indicates polarization as in Fig. 4. L. Example of four human-upregulated DA neuron DARs linked to human-specific upregulated DA neuron DEG NRN1. DARs, differentially accessible regions; FDR, false discovery rate; ns, not significant; hIns, human-specific insertions; hDels, human-specific deletions; HAQERs, human ancestor quickly evolved regions; ZooHARs, Zoonomia-defined human accelerated regions; hCONDELs, human-specific deletions in conserved regions; AADs, archaic ancestry deserts; DEGs, differential expressed genes

The majority of peaks called in human, chimpanzee, and macaque D40-100 cell types were located distal to gene promoters, in introns, and intergenic regions with a smaller proportion of peaks falling in exons and promoter regions (Figure 5B). After cross-species peak merging, we quantified read counts in each species and performed quality control (Figure S7A), which yielded 80,123 cells with high-quality paired snRNA-seq and snATAC-seq data (human = 23,962, chimpanzee = 33,016, macaque = 23,145). Integrating across species and performing dimensionality reduction on the snATAC-seq dataset while annotating cells with their snRNA-seq-defined cell type revealed similar structure in the snATAC-seq dataset as described above for snRNA-seq, with progenitors clustering separately from neuronal cell types and neurons separating by neurotransmitter and regional identity (Figures 5C, S7B, and S7C). Canonical marker genes for DA lineage cell types showed conserved promoter accessibility across species (Figure 5D), supporting the quality of the snATAC-seq dataset and cell type annotations.

To identify candidate *cis*-regulatory differences across species, we utilized the dreamlet package to perform differential accessibility testing of cross-species consensus peaks (species-only peaks meeting the same accessibility threshold were included separately in later analyses, Methods) across pseudobulk ATAC-seq samples within DA lineage cell types, adjusting our approach slightly compared to the RNA-seq model described above to improve modeling of inherently sparse ATAC-seq data (Methods). Most human-chimpanzee DARs that we identified were specific to progenitor, immature, or mature neuronal stages, with a smaller percentage shared across the DA lineage (Figure S7D). Focusing on DA neurons, human-chimpanzee differentially accessible regions (DARs) were predominantly located in noncoding regions of the genome and similar numbers of DARs had higher accessibility in each species (Figures 5E and 5F).

To study human-specific regulation in DA neurons, we intersected DARs and species-only peaks with human-specific evolutionary variants, including human-specific insertions (hIns)^51^, human-specific deletions (hDels)^51^, human ancestor quickly evolved regions (HAQERs)^52^, Zoonomia-defined human accelerated regions (zooHARs)^53^, indel-sized human-specific deletions in conserved regions (hCONDELs)^54^, and archaic hominin ancestry (admixture and incomplete lineage sorting) deserts (AADs)^55^, and found that hDels and hIns were significantly enriched in human-chimpanzee DARs (compared to a background set of all accessible peaks included in the analysis) (Figure 5G and Table S5), emphasizing the contribution of structural variants to *cis*-regulatory evolution.

To understand categories of genes that could be regulated by DARs, we annotated DARs using GREAT^56,57^. A majority of the top gene ontology (GO) terms were related to cyclic nucleotide biosynthesis and metabolism, including cyclic guanosine monophosphate (cGMP) signaling (Figures S7J and S7L). DARs within the regulatory domains of related genes were split between those with higher chromatin accessibility in human versus chimpanzee (42% and 58% of 50 DARs, respectively). Only 18% of these genes were differentially expressed between human and chimpanzee (46% of genes with above-threshold expression). Since cyclic nucleotide signaling is context-dependent, changes in gene regulation may only become apparent at the gene expression level in response to specific environmental or cellular stimuli. Interestingly, increased cGMP levels promote DA neuron mitochondrial function and may be neuroprotective in mouse models of PD^58^.

To further explore the concordance between differential accessibility and gene expression, we linked accessible chromatin regions to genes in both human and chimpanzee using Cicero, which utilizes a regularized correlation metric to predict co-accessibility in snATAC-seq data^59^. Most peaks were linked to the same gene in both species, with a smaller percentage linked to different genes or linked only in one species (Figure 5H). Each gene had a median of 6 linked peaks, and the majority of peaks were linked to a single gene in both species (Figures S7E and S7F). Since the total number of peak-gene links depended on the threshold applied to co-accessibility scores between accessible regions (with higher thresholds retaining only links with the strongest correlations), we plotted results across thresholds (Figure 5I). The percent of DE genes with linked DARs (regardless of direction) ranged from 6% at the most stringent threshold to 40% at a threshold of 0 (retains all links between regions within 500 kb), within the range of previous reports^50^. Reasoning that higher signal-to-noise would allow more reliable identification of correlations between regions with higher accessibility, we defined percent concordance as the percent of DARs with increased accessibility linked to upregulated DEGs, considering links defined in the species where the gene was upregulated. The percent concordance ranged from near-chance level (54%) at a threshold of 0 to 74% at the most stringent threshold (Figure 5I). At an intermediate threshold, approximately two thirds of DARs linked to upregulated DEGs in each species had increased accessibility in that species (Figure 5J, S7G, and S7H).

Although the majority of DARs were located distally with respect to gene promoters, DARs that overlapped promoters of DEGs had a high level of concordance with gene expression (80% of DAR-DEG promoter pairs representing 72 peaks and 61 genes). A representative example is RNAscope-validated human-upregulated gene *KCNJ16*, whose promoter overlaps two DARs with higher accessibility in human compared to chimpanzee and macaque (Figure S7K).

We next investigated DARs that overlapped enriched categories of human-specific sequence variation (Figure 5G). One prominent example is a DAR located about 2.5 kb upstream of the *GABRG3* promoter which partially overlaps a 326-bp human-specific insertion. This DAR had reduced accessibility in human compared to chimpanzee (p = 0.009) and *GABRG3*, which encodes an isoform of the gamma subunit of the GABAA receptor, displayed human-specific downregulation (p = 1.39e-06) (Figure 5K).

We also ranked genes by the number of concordant linked DARs. Interestingly, several of the top genes are known or predicted to be directly involved in neurite projection organization, including *EPHA10*, *NRN1*, and *NTNG2* (minimum ranks 2, 7, and 7, respectively), and all of these genes were upregulated in human compared to chimpanzee (*EPHA10*: p = 1.14e-19, *NRN1*: p = 5.24e-06, *NTNG2*: p = 4.72e-10). *NRN1*, for example, was classified as a human-specific upregulated gene and had three linked DARs (one intronic, p = 0.008 and two intergenic with p = 0.023 and p = 0.062) with increased accessibility in human, as well as slightly but not significantly increased promoter accessibility (Figure 5L). The *NRN1* gene encodes a small, extracellular cell surface protein which has increased expression in response to neuronal activity^60^, promotes axon outgrowth in retinal ganglion cells^61^ and has been shown to be neuroprotective in neurodegenerative disease^62,63^.

In summary, we present a resource of regions with differential chromatin accessibility between human and chimpanzee across the DA lineage, identify structural variations that may underlie species differences, and link DARs to candidate genes whose expression they may regulate.

### Conserved and divergent gene regulatory networks in ventral midbrain specification and maturation

We next examined the regulatory network context for species differences in gene expression and accessibility. Using SCENIC+^64^, we identified enhancer-driven gene regulatory networks (eGRNs) throughout the specification and maturation of midbrain cell types (Figures 6A-6C, S7P and S7Q, Methods). At D16, the analysis recovered 113 TF-driven eGRNs in human and 91 in chimpanzee, with a median size of 82 regions and 49.5 genes (Table S6). These eGRNs include many conserved master regulators with established roles in conferring or maintaining regional identity. These hub TFs were enriched in the expected progenitor subtypes, including *LMX1A*, *EN1* and *FOXA1* in midbrain progenitors (Figures S7P and S7Q). In addition, TFs with important roles for progenitor to neuron specification, such as *SOX2*, *HES1*, *POU3F2*, were active at different stages of maturation (Figures S7P and S7Q).

**Figure 6:**
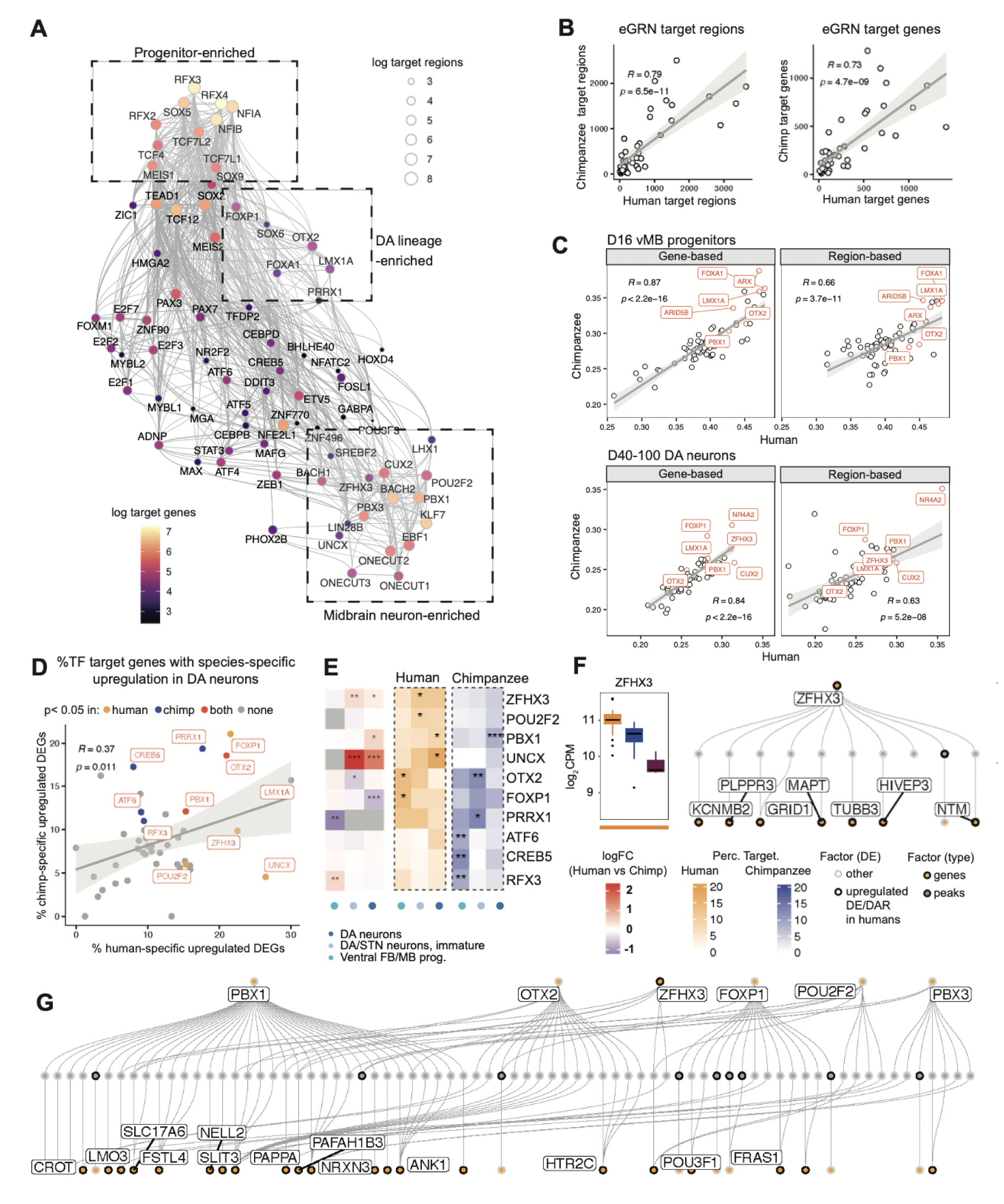
Conserved and divergent gene regulatory networks in ventral midbrain specification and maturation. A. Activator eGRNs in the developing human midbrain organoids (D40-100) projected in a weighted UMAP based on co-expression and co-regulatory patterns. Nodes label hub TFs of eGRNs with number of target genes (color) and target regions (size) plotted, and edges label co-regulatory networks between eGRNs. B. Scatterplots of number of genes (right) and number of regions (left) in activator eGRNs identified in humans versus chimpanzees. C. Scatterplots of eGRN specificity score in D16 vMB progenitors and D40-100 DA neurons in human versus chimpanzee, calculated based on target gene expression (left) and target region accessibility (right). D. eGRNs that have targets enriched for species-specific upregulated DEGs against other DEGs in DA lineage in humans or chimpanzees identified by Fisher’s exact test (p< 0.05, where other DEGs are defined as DEGs that are either not human/chimpanzee specific, or human/chimpanzee specific but down regulated in human/chimpanzee). Unions of DEGs from 3 D40-100 cell types (ventral FB/MB progenitors, immature DA/STN neurons, and DA neurons) in the DA lineage were used for testing. TFs are colored if corresponding eGRN targets are significantly enriched for human (yellow), chimpanzee (blue) or both (red) species-specific up-regulated DEGs. E. Heatmaps for differential expression of hub TFs showing log2FC in human versus chimpanzee (left), and for percentage overlap of species-specific upregulated DEGs in eGRNs in human (middle) and chimpanzee (right) DA lineage cell types. P values were calculated using Fisher’s exact test against overlap with other DEGs. F. Boxplot of *ZFHX3* expression across species in immature DA/STN neurons (line at the bottom represents human-specific polarization category) and human *ZFHX3* eGRNs (formulated as TF-peaks-genes) intersecting with upregulated DEGs or DARs in human immature DA/STN neurons. G. Human eGRNs enriched for upregulated DEGs or DARs in human DA neurons from Fig 6E. Nodes were pruned to include only the top 1500 highly variable genes and peaks and nodes connected to DEGs and/or DARs are shown. Genes or peaks that are human-specific upregulated in DA neurons are highlighted with a black border and genes that are connected with the most edges are labeled. eGRN, enhancer-driven gene regulatory network; TF, transcription factor; CPM, counts per million; DEGs, differential expressed genes; DARs, differentially accessible regions. * p < 0.05, ** p < 0.01, *** p < 0.001

In D40-100 organoids, 84 human and 114 chimpanzee eGRNs were recovered with a median size of 115 regions and 75.5 genes (Table S6). Since repressive interactions are more challenging to predict^64^, we chose to focus only on activator networks for downstream analyses. Visualizing human eGRNs based on TF coexpression and co-regulatory patterns highlighted modules that are enriched in midbrain progenitors, neurons, and the DA lineage (Figure 6A). These included eGRNs driven by *PBX1* and *NR4A2* (Figures 6A and 6C), genes with known functions for the development and survival of DA neurons and with disrupted functions in PD^65,66^. The numbers of regions and target genes per eGRN were correlated between species (Figure 6B). Similarly, focusing on the DA lineage, the specificity scores of networks were also correlated between species both when calculated based on target region accessibility or gene expression (Figure 6C), supporting broad conservation of *trans-*regulatory network membership and specificity.

We next asked whether individual eGRNs were enriched in species-specific upregulated genes in the DA lineage in human and chimpanzee, focusing on transcriptional activators identified in both species. Overall, we found a correlation in the fraction of species-specific upregulated DEGs within human and chimpanzee eGRNs (Figure 6D). Considering individual cell types, *OTX2*- and *PBX1*-driven networks were enriched for containing upregulated genes in both species, consistent with these eGRNs representing a substrate for regulatory alteration in each lineage. In contrast, some eGRNs, such as those driven by *PRRX1*, *POU2F2*, *UNCX* and *ZFHX3* were enriched for upregulated genes in only one species (Figure 6E), with *POU2F2* also enriched for upregulated DARs in humans (Figure S7M). Although the *UNCX*-driven eGRN was enriched for upregulated DEGs in human DA neurons, *UNCX* itself was chimpanzee-specifically downregulated, suggesting a derived change in the chimpanzee lineage (Figure S7N). However, *ZFHX3* showed a derived upregulation in human immature DA neurons and its eGRN was enriched for upregulated target genes in human, including microtubule genes *MAPT* and *TUBB3*, representing a candidate human-specific *trans-*regulatory alteration (Figure 6F). Consistent with *trans*-regulatory changes downstream of *ZFHX3*, comparative studies of human and chimpanzee PSCs revealed an excess of species-specific binding sites^67^. Meanwhile, several genes upregulated in humans that are related to antioxidant activity, including *PRDX2*, *PRDX3*, *PRDX4*, and *PRDX5* (Figures 4H and 4I), were predicted to be downstream of *NFE2L1* (*NRF1*), a TF depleted in DA neurons from individuals with PD with an established role in mediating protective oxidative stress responses^66,68^ (Figure S7O). Together, these analyses indicate a mainly conserved *trans*-regulatory landscape during ape ventral midbrain development, while highlighting several DA lineage-specific eGRNs enriched in human-specific gene regulatory changes that overlap with DEGs and DARs (Figure 6G).

### Oxidative stress induces conserved and species-specific responses in DA neurons

Given the observation of human-specific differences in antioxidant-related gene expression during DA neuron development, we next investigated if induced oxidative stress^27,28^ could unmask further functionally relevant species differences. We used rotenone, a toxic pesticide associated with PD risk that interferes with the mitochondrial electron transport chain, creating reactive oxygen species^69^. We optimized the rotenone concentration using single-species human and chimpanzee organoids to obtain a robust transcriptional response in both species without complete loss of DA neurons (Figures S8A-S8D). We applied the optimal concentration to interspecies organoids (Figure 7A), and used immunohistochemistry to visualize the progressive loss of TH^+^ neurons and fibers (Figures 7B and 7C, S8E-S8J). We collected multiomic data after 24 and 72 hours and after quality and doublet filtering, 28,552 nuclei were retained (Figures S9A-S9D), of which 3,087 were classified as DA neurons (human=1,442, chimpanzee=1,645 cells) based on known marker gene expression (Figures S9E-S9G). An important caveat is that some degenerating DA neurons may not have been identified due to loss of subtype-specific marker expression during the stress response (Figure S9F).

**Figure 7:**
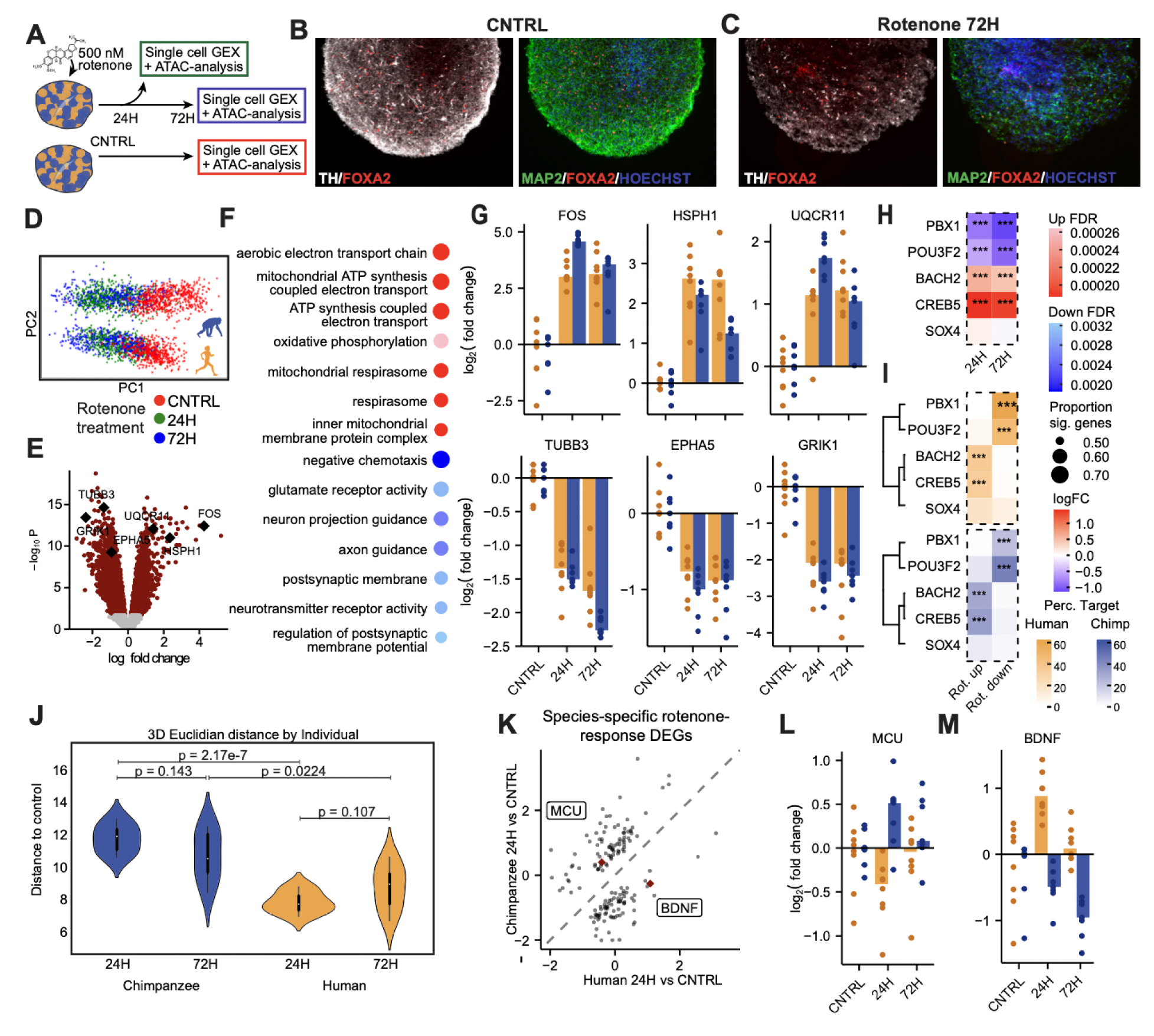
Oxidative stress induces conserved and species-specific responses in DA neurons in interspecies organoids. A. Experimental design for rotenone induced oxidative stress in interspecies organoids. D80 interspecies organoids were treated with 500 nM rotenone for 24- and 72 hours and were then immediately collected for multiomic snRNA- and ATAC-sequencing. B-C. Representative images of TH/FOXA2 and MAP2/FOXA2 immunohistochemistry in control organoids (B) and in organoids treated with rotenone for 72 hours (C), showing stress induced loss of TH^+^ cells and fibers. D. PCA plot after subsetting DA neurons, colored by condition with the bottom trajectory corresponding to human cells and the top trajectory corresponding to chimpanzee cells. E. Volcano plot of condition DEGs (average response in human and chimpanzee, FDR < 0.05) between control and 24 hours of rotenone treatment. F. Top GO terms for 24 hours of rotenone treatment versus control ranked by the proportion of genes with FDR < 0.05. G. Bar plots showing log_2_(fold change) for selected genes in each individual (dots) across the timecourse of rotenone treatment (normalized to control median expression across individuals within each species). H. Heatmap showing log_2_(fold change) for 24 hours and 72 hours of rotenone treatment (average response in human and chimpanzee) versus control. I. Heatmap for percentage overlap of eGRNs targets and upregulated or downregulated DEGs in human (top) and chimpanzee (bottom) under rotenone treatment in 24 and 72 hours in comparison to control. P values were calculated using Fisher’s exact test against overlap with other DEGs. J. Violin plots summarize the distribution of Euclidean distances in PCA space (PC1-3) from control to rotenone timepoint centroids across individuals. K. Scatterplot showing log_2_(fold change) between 24 hours of rotenone treatment and control in human versus chimpanzee for genes that were significant for the interaction of species and condition (FDR < 0.1) at 24 hours of rotenone treatment. L-M. Bar plots showing log_2_(fold change) across the timecourse of rotenone treatment (normalized to control within each species) for examples of genes that were significant for the interaction of species and condition at 24 hours of rotenone treatment: MCU (L) and BDNF (M). Scale bars: 200 μm. GEX, gene expression; ATAC, assay for transposase-accessible chromatin; FDR, false discovery rate; CNTRL, no rotenone; 24H, 24 hours of rotenone treatment; 72H, 72 hours of rotenone treatment, Rot., rotenone * p < 0.05, ** p < 0.01, *** p < 0.001

PCA of DA neuron transcriptomes revealed that responses to rotenone represented the major source of transcriptional variation along PC1 in both human and chimpanzee (Figure 7D), with early response genes, heatshock proteins, and cellular respiration genes marking increased rotenone responses (Figure S9H). Analysis of differential expression across the rotenone time course in both species using dreamlet revealed correlated (r = 0.87) responses at 24 and 72 hours (Figures 7E and S7I), enriched for GO terms associated with downregulation of neuronal functions such as axon guidance, neurotransmitter receptor activity, and synapses, consistent with previous studies^28^, and upregulated terms related to cellular respiration and ATP synthesis, possibly representing a compensatory response to electron transport chain inhibition^70^ (Figure 7F). Representative genes from the top categories showed generally conserved responses in human and chimpanzee (Figure 7G).

Next, we investigated candidate TFs driving gene expression shifts in response to oxidative stress in human and chimpanzee DA neurons. Using SCENIC+, we recovered 23 human and 13 chimpanzee eGRNs, with a median size of 51 regions and 47 genes (Table S6). Intersecting shared activator eGRNs between species with condition DEGs revealed a conserved *trans*-regulatory response upon rotenone treatment. Developmental TFs *PBX1* and *POU3F2* displayed reduced expression upon rotenone exposure at both time points and drove eGRNs enriched for downregulated genes in both species (Figures 7H and 7I). In addition, TFs *BACH2* and *CREB5*, both upregulated under oxidative stress, also served a conserved role in driving eGRNs that were enriched for upregulated genes following rotenone exposure (Figures 7H and 7I). These results suggest that eGRN analysis can provide insight into upstream regulators of condition-dependent transcriptional responses and highlight a conserved stress-induced gene regulatory landscape.

We further examined species-specific responses to rotenone using the species-condition interaction term in our model. Overall, there was broad conservation in oxidative stress response genes and regulatory networks, but human DA neurons had slightly blunted responses compared to chimpanzee neurons, in terms of both up- and down-regulated gene sets (Figures S9J and S9K). To further investigate the differences in the magnitude of rotenone induced transcriptional responses, we calculated the Euclidean distance in PC space and measured the impact of the stress response across individuals^30^(Figures 7J and S9L). Remarkably, all 7 chimpanzee individuals showed stronger responses across a similar trajectory than all 8 human individuals at both the 24 hour and 72 hour timepoints, when cultured together in the same interspecies organoid environment.

In addition, we identified dozens of genes with species-specific responses (Figure 7K, Table S7). Two examples of qualitative species differences include human-specific reduction of Mitochondrial Calcium Uniporter (*MCU*) (Figures 7L, S9M and S9N) and human induction of Brain-Derived Neurotrophic Factor (*BDNF*) (Figures 7M, S9O and S9P), both of which are consistent with increased neuroprotective mechanisms in human DA neurons. Notably, eGRN analysis suggested a possible species difference in *trans*-regulation of *BDNF* that could underlie divergent responses. While developmental TFs *PBX1*, *BACH1*, and *POU3F2* are the most highly ranked *BDNF* regulators in chimpanzee DA neurons, these TFs are ranked lower in human and instead *NR4A2* and early response TFs *JUNB* and *JUND* play more prominent roles (Figure S9Q).

Together, these findings reveal a conserved trajectory of oxidative stress response governed by shared eGRNs, while highlighting candidate genes with human-specific divergence that may have therapeutic implications.

## Discussion

The human DA system, even compared to that of our closest living ape relatives, more densely innervates larger target regions in the expanded human brain^71^. The molecular underpinnings driving these evolutionary changes and the consequences of the associated increase in bioenergetic demands remain unexplored. We hypothesized that the increased pressure on the human DA system may have necessitated adaptations in human DA neurons that could make them more resilient to cellular stress than the neurons of nonhuman primates in equivalent conditions. While these adaptations may be incomplete as suggested by increased human susceptibility to Parkinson’s disease over our longer lifespans, understanding potential protective mechanisms in human neurons could lay the foundation for future efforts to enhance or expand them with implications for disease treatment. Here, we established a phylogeny-in-a-dish approach, extending the concept of pooled iPSC culture systems^27,30,72^ by differentiating pooled midbrain cultures from 19 individuals across four primate species, facilitating the modeling of inter-versus intraspecies differences at baseline and in response to an oxidative stress challenge.

The gene expression landscape is highly dynamic during brain development, and the origin of connectivity differences between closely related primate species is unclear. To explore the extent and timing of human-specific gene expression patterns in DA neuron development, we sequenced progenitors and neurons from multiple timepoints, representing early stages of progenitor specification, neurogenesis and neuronal maturation. In immature DA neurons, the top categories of human-upregulated genes relate to the transport of mitochondria along axons/microtubules. This could reflect a strategy to meet the increased bioenergetic demands associated with the expanded axonal arborization of human DA neurons. Interestingly, *ZFHX3* also shows a quantitative human-specific upregulation during this transient window and acts as a hub TF for an eGRN enriched for genes quantitatively upregulated in human DA neurons including *MAPT*, *TUBB3*, and *NTM*. *ZFHX3* is expressed throughout the early developing brain and *ZFHX3* haploinsufficiency is associated with intellectual disability^73^. Although its function is not yet established in DA neurons, *ZFHX3* expression has previously been identified in a subset of high *TH/SLC6A3*-expressing DA neurons^74^*. ZFHX3* is involved in differentiation^75^, cytoskeletal organization^73^, survival, and protection against oxidative stress in other neuronal subtypes^75,76^, suggesting that increased *ZFHX3* expression may represent a consequential *trans*-regulatory change in developing human DA neurons. Additionally, several of the DEGs with the highest numbers of concordant linked DARs in DA neurons, including *NRN1*, are involved in neuron projection development. Together, these results reveal intriguing candidate genes that may be involved in promoting expanded DA neuron arborization and/or compensating for scaling-related consequences in the human brain.

The expanded innervation of human DA neurons likely increases metabolic demands related to the long-distance propagation of action potentials and the production of dopamine to supply the increased number of release sites^4,17^. Supporting the hypothesis that increased baseline oxidative stress is an important driver of gene expression divergence in humans, the top human-upregulated gene set in the most mature DA neurons in our dataset was related to antioxidant activity. This gene set included *CAT*, which encodes the hydrogen peroxide-degrading enzyme catalase. *CAT* was also found to be upregulated in human primary cortical tissue relative to chimpanzee and macaque^50^, supporting the relevance of our stem cell model to human evolution. In addition to catalase, genes encoding several members of the peroxiredoxin family of antioxidant enzymes, including *PRDX2*, *PRDX3*, *PRDX4*, and *PRDX5*, were also upregulated in human DA neurons. Interestingly, this gene family is markedly expanded and under positive selection in cetaceans, which has been suggested to protect against reactive oxygen species generated during diving-related hypoxia^77^. Both catalase and peroxidase activity have been found to be reduced in the brains of PD patients^78^. Overexpression of *PRDX2* and *PRDX5* decreases the toxicity of PD-inducing chemicals^79,80^, while silencing of *PRDX5* results in increased sensitivity to rotenone-induced death^81^. While further studies are required to examine the functional consequences of the increased expression of these genes, our results build on theoretical proposals^4,10^ by demonstrating human-specific mechanisms that could protect DA neurons from increased oxidative stress.

The finding that protective mechanisms against oxidative stress display human-specific quantitative increases at time points corresponding to fetal stages motivated us to investigate whether induced oxidative stress could unmask additional species differences. While the response to rotenone was mediated by conserved eGRNs, there were also several notable differences. We observed a significantly blunted transcriptional response to rotenone in humans in every individual in our panel. At the level of individual genes, the expression of the mitochondrial Ca^2+^ uniporter gene *MCU* was already lower in human at baseline, with expression further reduced upon rotenone exposure, while in chimpanzee the expression was increased. *BDNF* expression showed the opposite trend, with rotenone exposure resulting in reduced expression in chimpanzee but increased expression in human DA neurons. Both of these examples suggest an increase in neuroprotective mechanisms in human neurons. The deletion of *MCU* has been shown to be neuroprotective in both genetic and toxin-induced models of PD^82,83^, and MCU blockers prevent iron-induced mitochondrial dysfunction^84^. Neurotrophic BDNF is essential for the development of DA neurons, promotes DA neuron survival in animal models of PD and is reduced in the substantia nigra of patients with PD^85–87^. Indeed, delivery of neurotrophins, including GDNF and BDNF, is under study as a therapeutic option in PD^87^. However, it is still unclear whether rotenone-induced *BDNF* expression is truly human-specific since at least one study also found increased *BDNF* transcription but not protein levels in mouse DA neurons with chronic *in vivo* rotenone exposure^88^. Future studies will be needed to determine whether the altered transcriptional response to rotenone provides a functional neuroprotective effect in human DA neurons. Taken together, these observations provide additional support for the hypothesis that human DA neurons have mechanisms for enhanced buffering of oxidative stress, both at baseline and following perturbation

This study opens questions related to the functional implications of the molecular changes that we identify. How does the genetic manipulation of candidate genes or regulatory elements with human-specific expression or accessibility affect intrinsic features of DA neurons such as axonogenesis, firing rate, dopamine release, and response to oxidative stress? While the diversity of our iPSC-derived midbrain cultures allowed us to study cell type-specific patterns of gene expression and regulation, this heterogeneity poses challenges for functional interrogation. Future experiments with CRISPR-mediated knockdown or inactivation in isogenic lines will be needed to dissect the contributions of individual genes and regulatory elements, and multimodal readouts will be essential. Transplantation of DA neurons into rodent models, where endogenous innervation patterns and connectivity can be recapitulated^33,89^ could allow studies of cell-intrinsic species differences in arborization and candidate gene function in more mature DA neurons.

Since the divergence of human and chimpanzee from a common ancestor, the human brain has evolved in both size and function, and in the susceptibility to neurodegenerative and neuropsychiatric diseases^90^. Many of these disorders involve the DA system, either directly through a progressive degeneration of DA neurons (PD), or as part of DA signaling dysregulation (e.g. autism spectrum disorder^91^, schizophrenia^92^, bipolar disorder^93^). By gaining molecular knowledge of what sets the human DA system apart from that of our closest extant relatives, we can advance our understanding of the origins of human-enriched disorders, and identify new therapeutic targets and strategies for drug development. In addition, while stem cell based cell replacement therapy for PD is currently in clinical trials as a potential restorative treatment strategy, there are still open questions related to the long-term viability of the transplanted neurons^94^. Our dataset represents a resource of tolerated interspecies variation that could be mined for candidate genes whose manipulation could improve the innervation, function, and/or survival of DA neurons in cell replacement paradigms.

### Limitations of the study

While we used state-of-the-art protocols to pattern and mature midbrain neurons^31^, the cell types studied here are still relatively immature (corresponding to mid to late fetal stages). The DA neuron subtype identity is not yet fully refined at fetal timepoints^74,95^, and we do not yet see separation of subtype-specific markers between different single cell clusters^96^, preventing conclusions about subtype-specific differences. Furthermore, while rotenone exposure allows us to simulate conditions of oxidative stress that might occur in the aging brain, it is unclear how context-dependent responses to oxidative stress might differ in fetal versus aged neurons. Future studies could integrate strategies to accelerate neuronal maturation^44^ or simulate aging^97^ to improve the maturity and relevance of these cell populations to human neurodegenerative disease. While we established a clear correlation between species differences in RNAseq data from primary tissue and in our organoid model, the limited availability and quality of primary tissue samples made it difficult to validate these results with orthogonal methods.

We chose to maximize statistical power by combining cells from inter- and intraspecies pools in our analysis. While the majority of human and chimpanzee DA neurons (human: 64.0%, chimpanzee: 75.1%) were derived from interspecies pools and gene expression was highly consistent across pool types (Figure 2J, S5C-D), pointing to the contribution of cell-intrinsic factors, future work will be needed to clarify the relative contributions of cell-intrinsic and cell-extrinsic influences to the human-specific patterns of gene expression and regulation we have identified. In addition, glia were mostly lacking in our organoid models, and they may be an important source of cell-extrinsic species differences^98,99^ and influence disease susceptibility^100^. Finally, locally supplied dopamine in DA target structures could represent an additional compensatory mechanism beyond those discussed here that could offset the increased demand for dopamine secretion in the human brain^71^.

## Resources

### Lead contact

Further information and requests for resources and reagents should be directed to and will be fulfilled by the lead contact, Alex A. Pollen.

### Materials availability

This study did not generate new unique reagents.

### Data and code availability

● Sequencing data are deposited on GEO accession number GSE292438
● Processed data available in browsable format: https://cells.ucsc.edu/?ds=xsp-dopa-organoids
● Original code is publicly available and can be found here:

○ https://github.com/nkschaefer/cellbouncer
○ https://github.com/jenellewallace/CrossPeak
● Any additional information required to reanalyze the data reported in this paper is available from the lead contact upon request.

## Supporting information

MRI measurements of human and macaque brain volumes, related to Figure 1.

Batch information for differentiation and sequencing, related to Figure 1 and 2

Differential gene expression analysis, related to Figure 4

Differential accessibility analysis, related to Figure 5

eGRN information in D16, D40-100, and Rotenone dataset, related to Figure 6 and 7

Condition- and species-dependent response to rotenone, related to Figure 7

Key Resource Table

## Acknowledgments

The authors thank Cynthia Dunbar and So Gun Hong for sharing of macaque iPSC lines, Danny Conrad and the Gartner lab for sharing MULTIseq reagents and for assistance in optimizing barcode labeling and A. Tarantal at UC Davis for providing samples. We thank Amanda Everitt and Katie Pollard for helpful discussions about cross-species differential expression and accessibility analysis and all members of the Pollen lab for invaluable feedback during the course of the work. Sequencing was performed at the UCSF CAT, supported by UCSF PBBR, RRP IMIA, and NIH 1S10OD028511-01 grants. RNAscope imaging was performed at Innovation Core at the Weill Institute for Neurosciences and organoid culture for HD-MEA recordings was done at the cell culture core facility at UCSC Institute for Biology of Stem Cells (RRID SCR_021353). This work was supported by the following funding sources:CIRM fellowship (SN), Schmidt Science Fellows (JLW), Jane Coffin Childs (JLW), NIH under award numbers K12GM139185 (JLS), 1RM1HG011543 (SRS) and R01MH120295 (SRS), UC Santa Cruz Chancellor’s Postdoctoral Fellowship (JLS). Lundbeck Foundation R380-2021-1267 (JK), Novo Nordisk Foundation NNF21CC0073729 (JK, AK), Weill Neurohub Fellowship (NKS), National Institutes of Health R01AG087959, 1R01MH134981, DP2MH122400 (AAP), NSF #2134955 (AAP), Pershing Square Foundation, Schmidt Futures Foundation, and the Shurl and Kay Curci Foundation Innovative Genomics Institute Award (AAP). AAP is a New York Stem Cell Foundation Robertson Investigator. This project was funded in part by the Emory National Primate Research Center Grant No. ORIP/OD P51OD011132.

## Author contributions

SN and AAP conceived the project, designed and supervised the experiments. SN generated pooled inter- and intra-species cultures and performed cell culture experiments. SN, JLW, MTS and AB prepared samples for 10x sequencing, and generated GEX and ATAC libraries. NKS developed and implemented methods for species- and individual demultiplexing. SN and JLW processed and clustered the single nuclei GEX data. JLW performed differential gene expression and accessibility analyses and developed methods for cross-species ATAC peak comparisons. JWD performed eGRN analyses,comparisons to the fetal data set and DA neuron maturation comparisons. RM performed cNMF simulations. MO performed analysis on the shift in euclidean distances following rotenone treatment. IB cultured organoids and collected samples for functional assessments, JLS performed and analyzed HD-MEA recordings and JK performed DA release experiments. EC and KH performed RNAscope experiments. SN performed immunochemistry experiments, with assistance from DS. BP provided and characterized human and chimpanzee iPSC-lines, and BP and DS performed whole genome sequencing on iPSC-lines. TZ and FW performed and analyzed MRI experiments. SN, JLW and JWD prepared the figures. SN, JLW and AAP wrote the paper with input from all authors. SRS, AK, HH, NKS and AAP supervised the work.

## Supplement figures

**Figure S1 (related to Figure 1 and 2):**
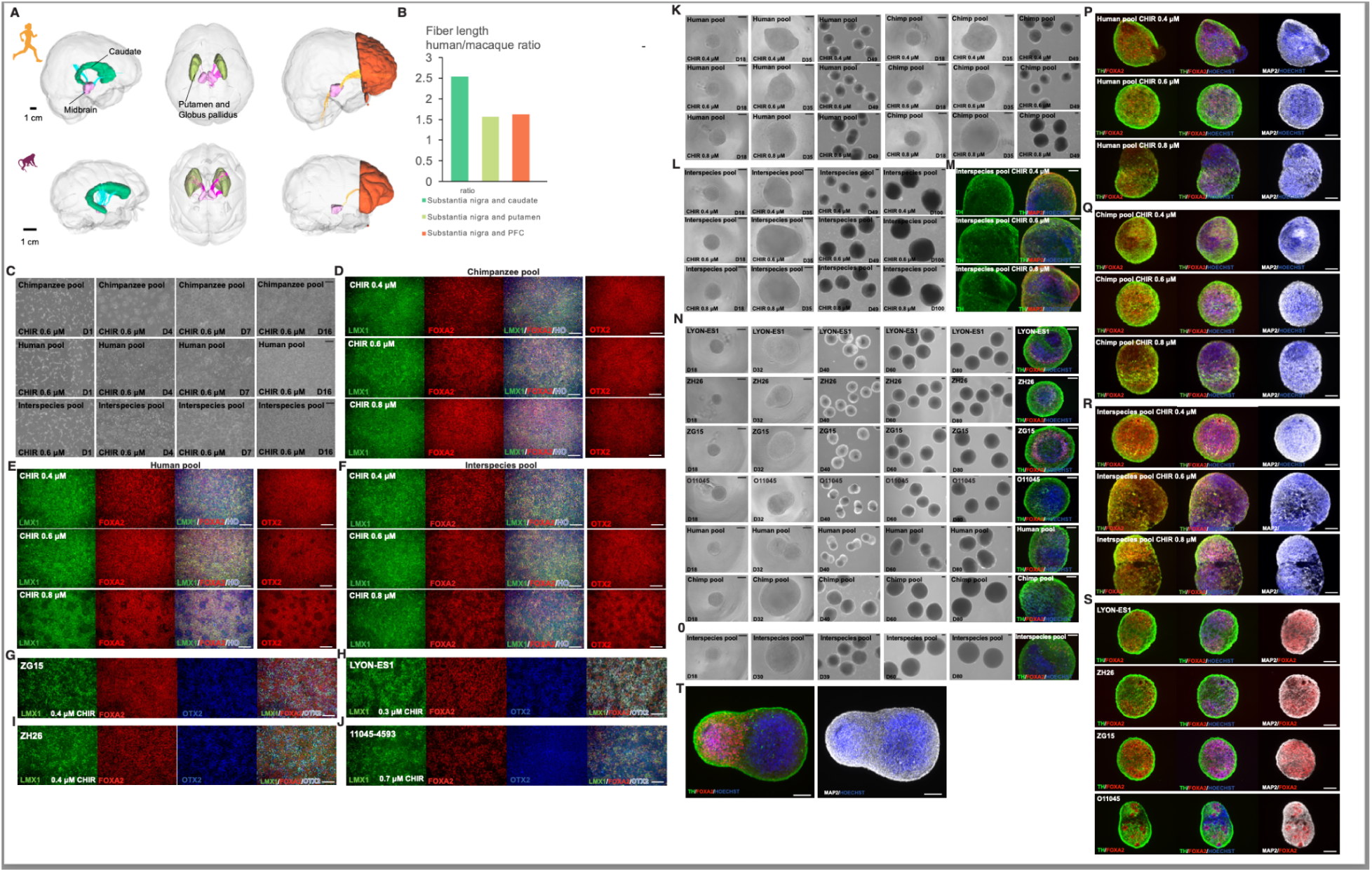
Generation and characterization of ventral midbrain intra- and interspecies progenitor pools and organoids from four primate species. A. Images highlighting human and macaque midbrains, dopamine target regions and dopamine fiber tracts that were measured through diffusion MRI to obtain ratios between species. B. Dopaminergic fiber length ratios between human and macaque (n=1 individual per species), but without taking into account differences in arborization within target regions. C. Brightfield images of inter- and intraspecies pools from D1 until D16 when the cells were dissociated and re-seeded for 3D culture, either before or after cryopreservation. D-F. Immunocytochemical characterization of chimpanzee pools (D), human pools (E) and interspecies pools (F) on D14, from three different CHIR concentrations. G-J. Immunocytochemical characterization of the four outgroup lines; ZG15 (G); LYON-ES1 (H); ZH26 (I); and 11045-4593 (J), that were differentiated individually in a subsequent experiment, in order to increase the number of macaque and orangutan cells and individuals. K. Brightfield images of organoids from intraspecies human or chimpanzee pools after seeding D16 progenitors for maturation in 3D. L. Brightfield images of organoids from interspecies pools after seeding D16 progenitors for maturation in 3D. M. D100 interspecies organoids across three CHIR concentrations, immunohistochemically labeled for TH, MAP2 and Hoechst. N. Brightfield images of organoids from the outgroup experiment, and including organoids from the intraspecies pools (CHIR 0.8 mM), and immunohistochemistry for TH/FOXA2 on D80. To generate the organoids, D16 progenitors were cryopreserved and then seeded directly into ultra-low attachment plates post thawing. O. Brightfield and immunohistochemistry (TH/FOXA2) images of interspecies organoids from the rotenone experiment (CHIR 0.8 uM), generated from cryopreserved D16 progenitors. P-R. Immunohistochemistry of human (P), chimpanzee (Q) and interspecies (R) pooled organoids on D40, from three different CHIR concentrations. S. Immunohistochemistry of macaque (LYON-ES1, ZG15, ZH26) and orangutan (O11045) individuals on D40. T. Immunohistochemistry of human midbrain - human cortical organoids, 6 days post-fusion on D40, with TH^+^ fibers being visible in the midbrain organoid and on the surface of the cortical organoid. Scale bars: 400 μm (C); 200 μm (D-T). PFC, prefrontal cortex; CHIR, CHIR99021.

**Figure S2 (related to Figure 1 and 2):**
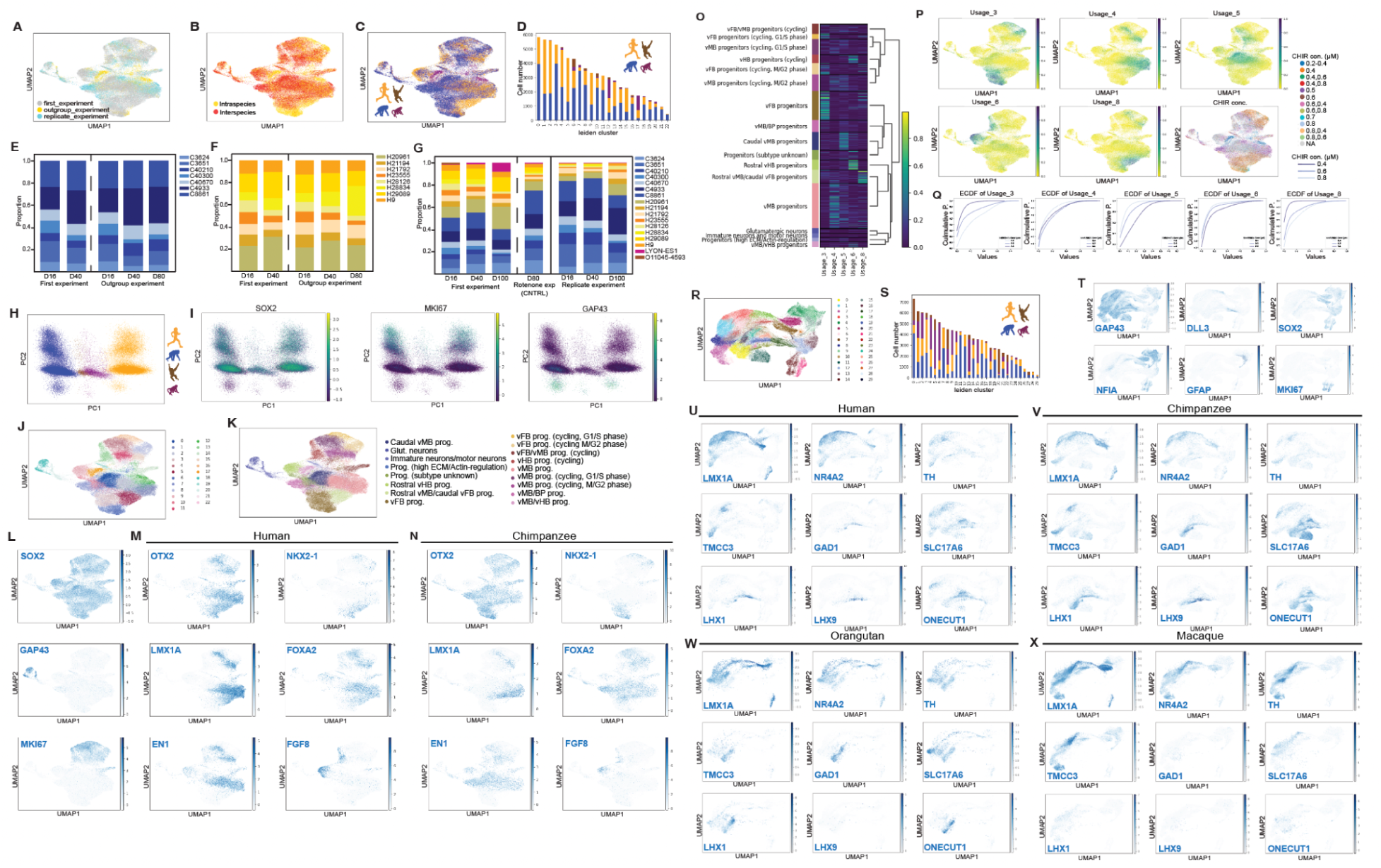
Species-, individual- and cell type composition of midbrain progenitor pools and intra- and interspecies organoids. A-C. UMAPs of D16 cells colored by experiment (A), pool type (B), and species (C). D. Bar chart showing the number of cells per species for each leiden cluster. E-G. Proportions of individuals in chimpanzee (E) and human (F) intraspecies pools, and interspecies pools (G), post quality control, doublet removal and removal of H28834 at D100. The dashed line separates pools that are derived from the same D16 progenitors, and the full line separates pools that were generated from a replicate pool, where the unequal proportions of human and chimpanzee were observed from the onset of differentiation. H. Principal component analysis (PCA) plot of D16 cells showing separation based on species identity along PC1. I. PCA plots of D16 cells with projection of *SOX2*, *MKI67* and *GAP43* expression levels to illustrate maturation and cell cycle state as major drivers of the separation along PC2. J-K. UMAPs of D16 cells colored by leiden (J) and assigned cell type (K). L. UMAPs of D16 cells colored by scaled and normalized expressions of *SOX2*, *MKI67* and *GAP43*. M-N. UMAPs of D16 cells from human (M) and chimpanzee (N) colored by scaled and normalized expression of genes with distinct regional distributions along the rostro-caudal axis. O. Heatmap showing usage of gene expression programs (GEPs) 3, 4, 5, 6, 8 in each cell type at D16. These GEPs showed distinct usage patterns corresponding to subtype identities. P. UMAPs of day 16 cells colored by usage of GEPs that correspond to subtype identities, or the applied CHIR concentration (lower right corner). Q. Empirical cumulative distribution plots of the usage of the cell type specific GEPs across CHIR concentrations at D16. High CHIR concentrations are associated with increased usage of GEPs that are used by posterior progenitor subtypes, while lower CHIR concentrations are associated with increased usage of GEPs that are used by anterior progenitor subtypes. R. UMAP of D40-100 cells, colored by leiden cluster. S. Bar chart of D40-100 cells, showing the number of cells per species for each leiden cluster. T. UMAPs of D40-100 cells, colored by scaled and normalized expression of neural stage and state specific genes. U-X. UMAPs of D40-100 cells, colored by scaled and normalized expression of neuronal subtype markers in human (U), chimpanzee (V), orangutan (W) and macaque (X). Prog, progenitors; vMB, ventral midbrain; vFB, ventral forebrain, vHB, ventral hindbrain; BP, basal plate; ECM, extracellular matrix; glut, glutamatergic; DA, dopaminergic; P, Probability

**Figure S3 (related to Figures 2 and 3):**
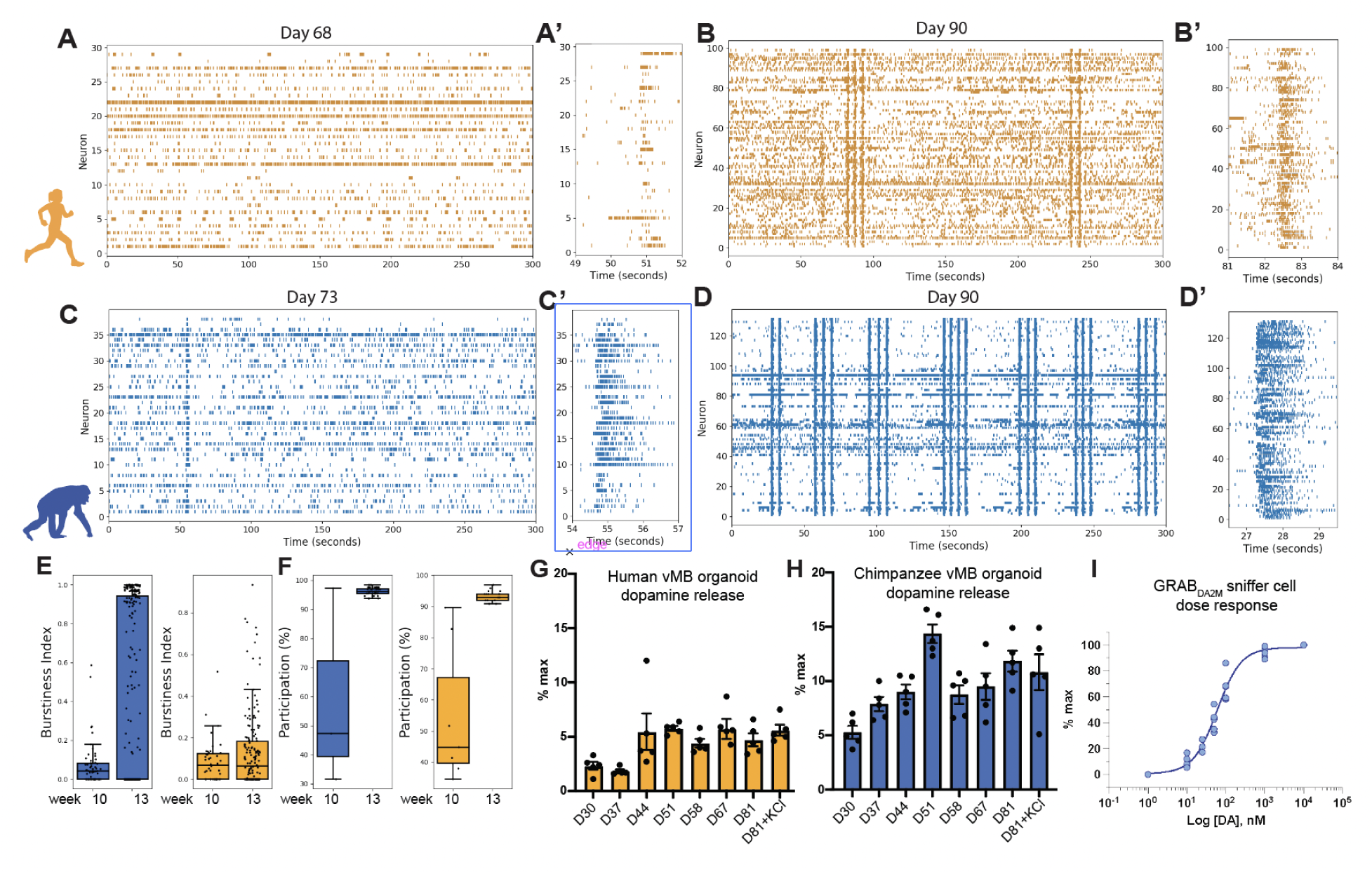
Functional characterizations of human and chimpanzee ventral midbrain organoids. A. Raster plot of endogenous spiking activity (orange dashes) in a D68 human-derived midbrain organoid measured from the 29 spike-sorted units, with a closeup on the first correlated network event (burst) from the recording in (A’). B. Raster plot of endogenous spiking activity (orange dashes) in the same human organoid at D90. Activity was detected in 99 spike-sorted units, with a closeup on the first correlated network event (burst) from the recording in (B’). C. Raster plot of endogenous spiking activity (blue dashes) in a D73 chimpanzee-derived midbrain organoid measured from the 38 spike-sorted units, with a closeup on the first correlated network event (burst) from the recording in (C’). D. Raster plot of endogenous spiking activity (blue dashes) in the same organoid at D90. Activity was detected in 131 spike-sorted units, with a closeup on the first correlated network event (burst) from the recording in (D’). E. Box-and-whisker plot showing burstiness index (defined as the number of spikes produced by a neuron within a bursting epoch / total number of spikes) at week 10 and week 13 in human- and chimpanzee-derived organoids. In chimpanzee organoids, burstiness increased significantly between week 10 and week 13 (p < 0.0001, Cohen’s d = -0.80), indicating a moderate effect. In human organoids, the increase in burstiness between the same time points was smaller but still statistically significant (p = 0.037, Cohen’s d = -0.26). Statistical comparisons were done using the Welch’s t-test, due to differences in sample sizes. F. Neuronal participation (defined as the percent of neurons with a firing rate greater than average during a bursting epoch) at week 10 and week 13 is shown. In chimpanzee organoids, the difference in participation between these timepoints was not statistically significant (p = 0.197), but it had a large effect size (Cohen’s d = -5.15) indicating that statistical power was limited by the relatively low number of bursts at the earlier time point. In human organoids, the increase in participation between week 10 and week 13 was statistically significant (p = 0.004, Cohen’s d = -3.27), and showed a very large effect. Statistical comparisons were done using the Welch’s t-test, due to differences in sample sizes. G-H. The spontaneous (D30-81) and KCl depolarization-induced (D81) secretion of dopamine was measured from human (G) and chimpanzee (H) intraspecies pooled organoids over time. I. Response curve of GRAB_DA2M_ showing the correlation between %max that is reported in (G-H) and the actual DA concentration in nM.

**Figure S4 (related to Figure 1 and 2):**
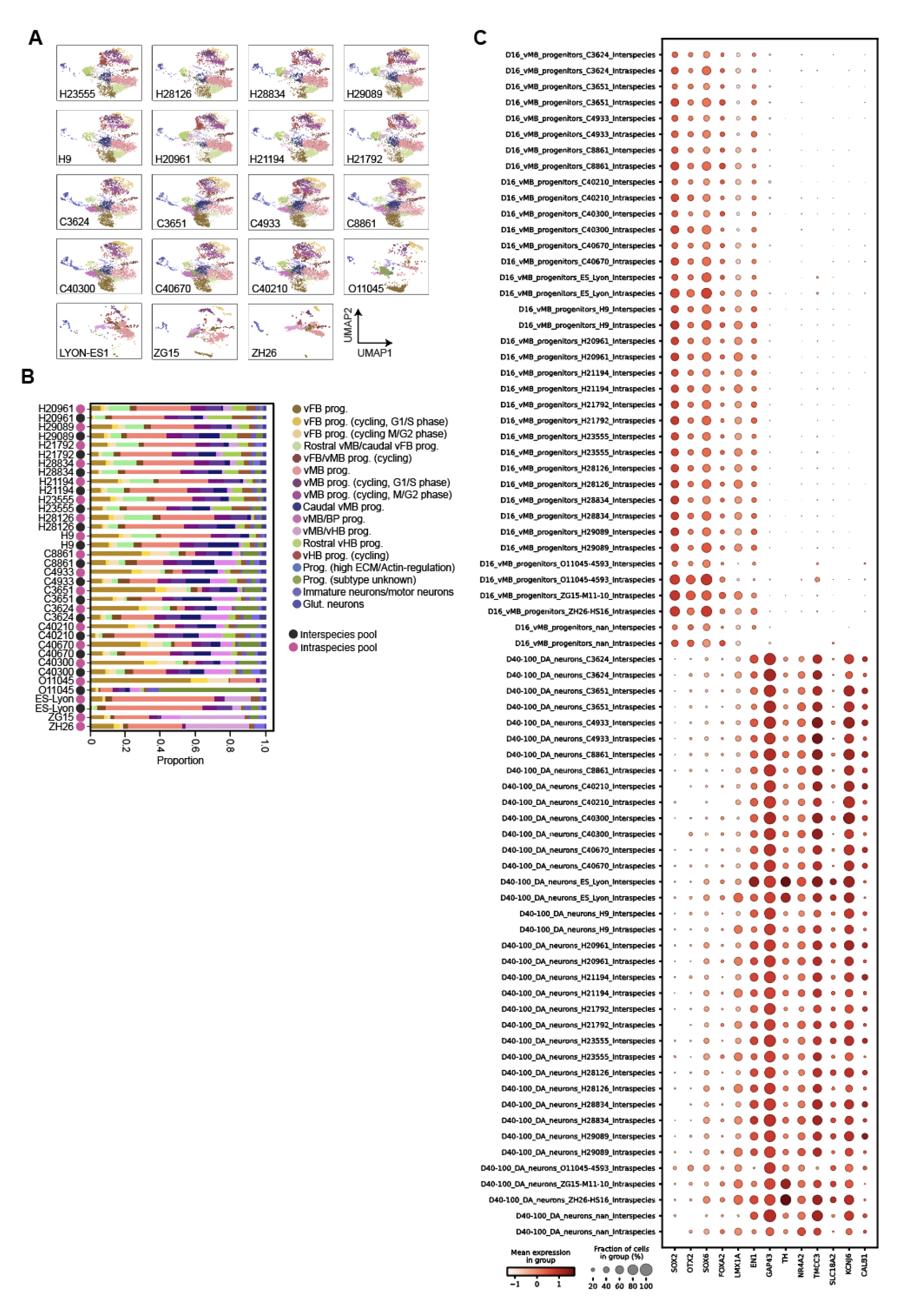
Conservation of cell type composition and DA lineage markers across individuals and pool types. A. UMAPs of D16 cells colored by cell type for each human, chimpanzee, orangutan and macaque individual. B. Stacked bar plots highlight that every human and chimpanzee individual is represented across all cell types in both pool types at day D16. C. Dotplots highlight marker gene expression levels and frequency across DA lineage cell types as separated by individual and pool type, highlighting the reproducibility of vMB progenitor and DA neuron specification across experimental conditions.

**Figure S5 (related to Figure 3 and 4):**
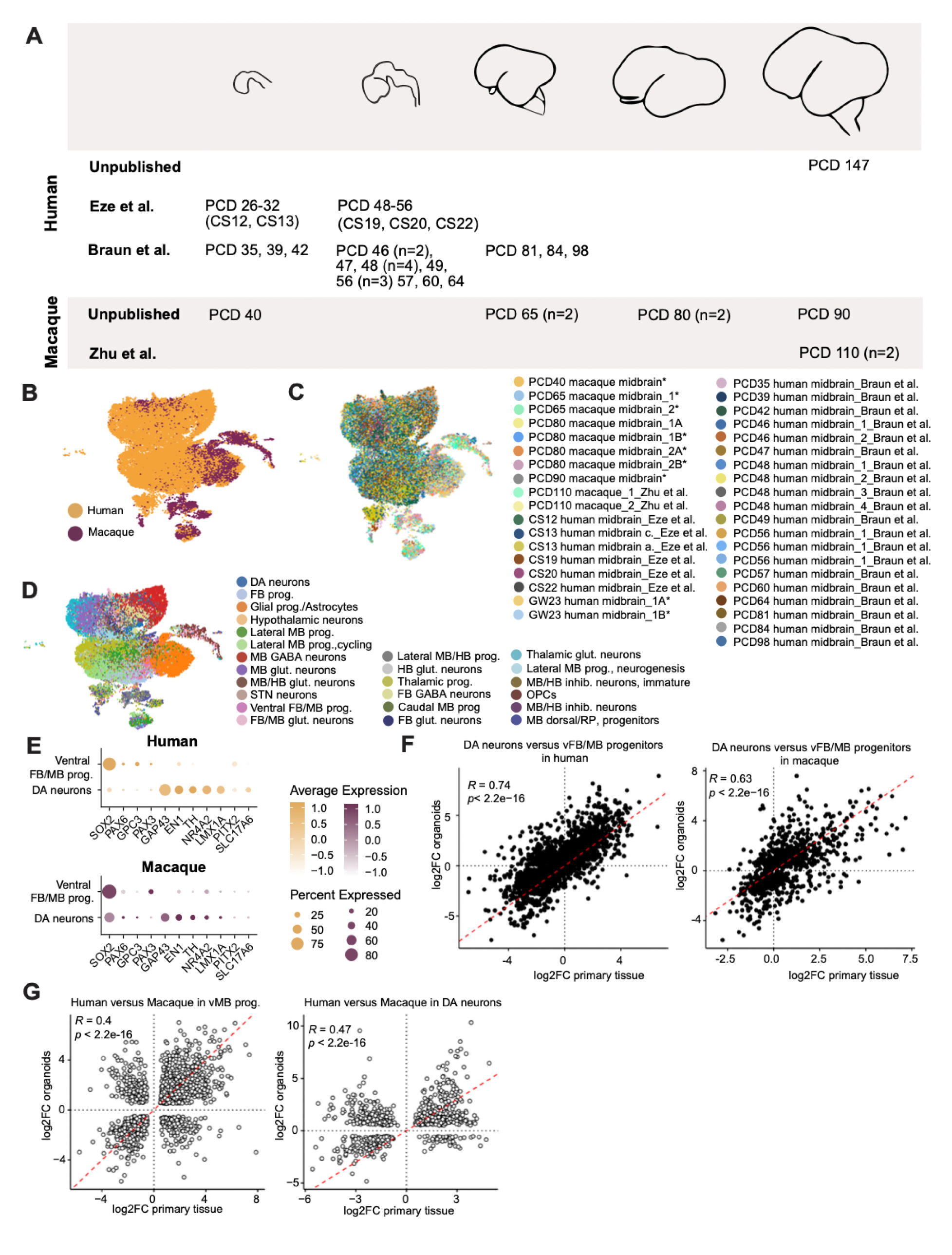
Human and macaque primary midbrain samples for cross-species comparisons of genes expression divergence. A. Overview of human and macaque primary midbrain samples that were included in the analysis, including one new human and 6 new macaque individuals collected in this study. B-D. UMAPs of primary midbrain cells colored by species (B), sample (* indicates new data collected in this study) (C), and cell type (D). E. Dotplots highlight marker gene expression levels and frequency in ventral FB/MB and progenitors and DA neurons across human (top) and macaque (bottom). F. Scatter plots for correlation of differentially expressed genes between vMB progenitors and DA neurons in vivo (primary) and in vitro (organoids) in human (left) and macaque (right). G. Scatter plot showing the log fold change of gene expression between orthologous human and rhesus macaque vMB progenitors (left) and DA neurons (right) for genes with significant differential expression in both primary and organoid datasets. In both cell types, a significant and positive correlation between fold changes across species is identified. Note that the analyses in F and G likely represent a lower bound on the correlation between primary and organoid differential expression because the primary datasets used individual species genomes for alignment and whole cell capture whereas for our organoid dataset, we used an ancestral *Homo*/*Pan* genome and nuclei capture.

**Figure S6 (related to Figure 4):**
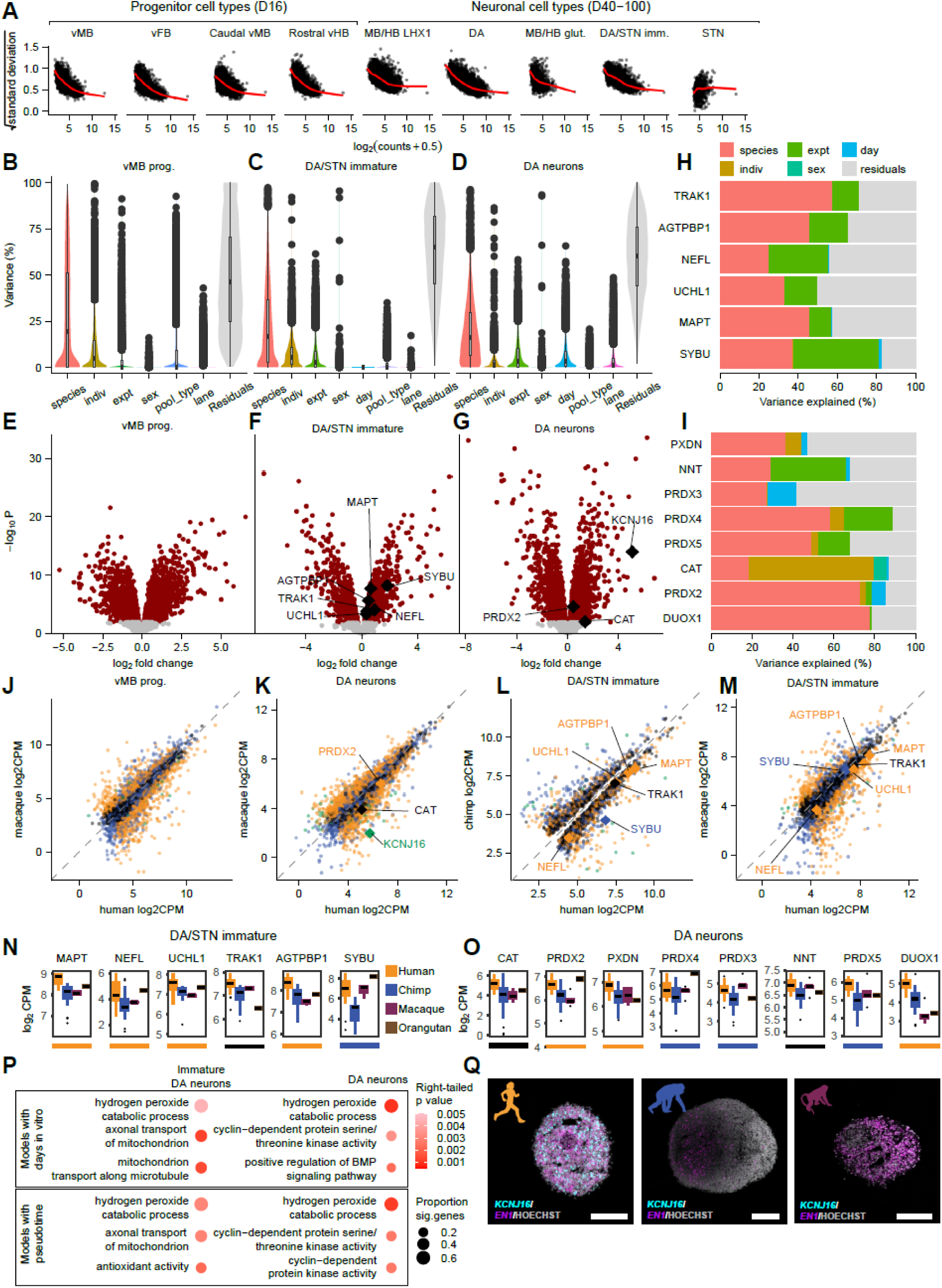
Modeling and validation of gene expression divergence across human and non-human primate ventral midbrain development. A. Voom plot showing the relationship between the mean count and standard deviation across pseudobulk samples for each gene. B. Variance partition plot for the full model including all tested variables for D16 vMB progenitors. C. Variance partition plot for the full model including all tested variables for D40-100 immature DA/STN neurons. D. Variance partition plot for the full model including all tested variables for D40-100 DA neurons. E. Volcano plot showing human-chimpanzee DEGs for D16 vMB progenitors. F. Volcano plot showing human-chip DEGs for D40-100 immature DA/STN neurons (genes from Fig 4G are labeled). G. Volcano plot showing human-chip DEGs for D40-100 DA neurons (genes from Fig 4I are labeled). H. Variance partition for genes shown in Figure 4G in immature DA/STN neurons (including only variables used in the final model from Figure 4). I. Variance partition for genes shown in Figure 4I in DA neurons (including only variables used in the final model from Figure 4). J. Scatterplot showing average normalized expression across pseudobulk samples for each human-chimpanzee DEG in human versus macaque D16 vMB progenitors, with points colored by categories from Figure 4C. K. Same as J but for D40-100 DA neurons. L. Same as J but for human versus chimpanzee in D40-100 immature DA/STN neurons. M. Same as L but for human versus macaque. N-O. Boxplots for human-chimpanzee DEGs belonging to the top human-upregulated GO term in immature DA/STN neurons (N) and DA neurons (O) showing normalized median gene expression values across species (human, chimpanzee, and macaque repeated from Figure 4, orangutan added) with colored lines below indicating polarization category. Note that orangutan expression levels were not considered when assigning polarization categories since we only included a single orangutan individual. P. Dotplot showing the top three upregulated GO terms in immature DA/STN neurons (left) and DA neurons (right), with dot size corresponding to the proportion of genes in the gene set that were significantly differentially expressed and color showing the unadjusted p values for a one-tailed test of gene set upregulation in human compared to chimpanzee. The top row shows results for the model described in Figure 4 where days in vitro was included as a continuous covariate in the linear mixed model and the bottom row shows an alternate model substituting Monocle3 pseudotime values. Q. RNAscope of *EN1* and *KCNJ16* in D40 organoids from human (left), chimpanzee (middle) and macaque (right). Scale bars: 200 μm (Q). Indiv = individual (cell line), expt = experiment

**Figure S7 (related to Figures 5 and 6):**
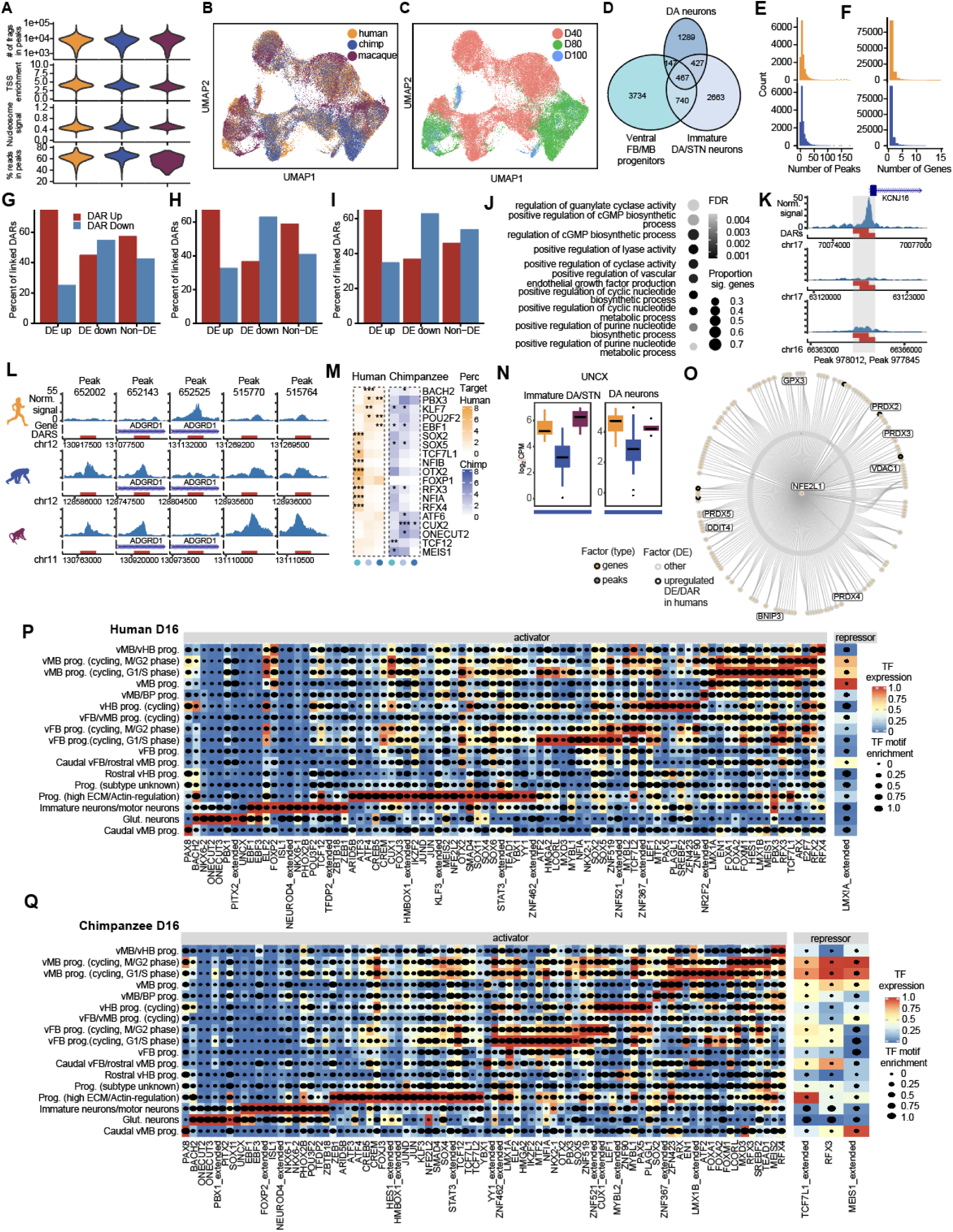
Evolution of the *cis*-regulatory landscape and regulatory networks in ventral midbrain specification and maturation. A. Violin plots showing distribution of quality scores in snATAC-seq data (based on peaks called on each species’ own genome before cross-species consensus peak merging) across human, chimpanzee, and macaque, calculated with Signac. Row by row from the top: number of fragments in peaks, transcription start site (TSS) enrichment, nucleosome signal, percent reads in peaks. B-C. Same UMAP as in Figure 5C but with species (B) and time point (C) labels. D. Area-proportional Venn diagram showing intersections between lists of human-chimpanzee DARs in DA lineage cell types. E. Histograms showing the number of peaks linked to each gene in DA lineage cell types with Cicero for human (top, median = 6 peaks) and chimpanzee (bottom, median = 6 peaks) F. Histograms showing the number of genes linked to each peak in DA lineage cell types with Cicero for human (top, median = 1 gene) and chimpanzee (bottom, median = 1 gene) G. Barplot showing the percent of DARs linked to DE up, DE down, and non-DE genes with Cicero that have increased versus decreased accessibility in human at co-accessibility threshold 0.15 (n = 181 human-up DEGs with 229 linked DARs, 171 human-down DEGs with 237 linked DARs, 280 non-DE genes with 355 linked DARs). H. Same as G but for chimpanzee. (n = 173 chimpanzee-up DEGswith 231 linked DARs, 198 chimpanzee-down DEGs with 265 linked DARs, 319 non-DEGs with 380 linked DARs). I. Same as G but for GREAT in human (n = 290 human-up DEGs with 414 nearby DARs, 318 human-down DEGs with 473 nearby DARs, 499 non-DEGs with 705 nearby DARs). J. Top 10 GO terms from GREAT analysis (foreground = DARs and human-only peaks, background = all peaks tested in dreamlet model and human-only peaks) ranked by FDR. Shading shows FDR and size shows the proportion of genes in the set that had a linked DAR. K. DARs overlapping the promoter region for *KCNJ16* in human (top), chimpanzee (middle), and macaque (bottom) in DA neurons. Due to inconsistent annotations for this gene across species, the human NCBI Refseq annotation is shown. L. DARs linked (via GREAT) to the *ADGRD1* gene (included in the top-ranked term from J) showing signal in DA neurons in human (top), chimpanzee (middle), and macaque (bottom). M. Heatmap for percentage overlap of human or chimpanzee upregulated DARs in eGRNs in DA lineage cell types. P values were calculated using Fisher’s exact test against overlap with other DARs, including those downregulated in the corresponding species or upregulated in other species (* p < 0.05, ** p < 0.01, *** p < 0.001) N. Boxplot of *UNCX* expression across species in human immature DA/STN neurons (left) and DA neurons (right) (lines at the bottom represents chimpanzee-specific polarization category). O. Human *NFE2L1* eGRN in the D40-100 dataset. Nodes are pruned to top highly variable genes or regions and genes under GO term ‘hydrogen peroxide catabolic process’ were labeled. Genes or peaks that are human specific upregulated in DA neurons were highlighted by black border P-Q. Heatmaps for shared D16 eGRNs between human (M) and chimpanzee (N). Color represents TF expression and size represents TF motif enrichment under each cell type. DARs, differentially accessible regions; DE, differential expression; FDR, false discovery rate; eGRN, enhancer-driven gene regulatory network; TF, transcription factor; CPM, counts per million.

**Figure S8 (related to Figure 7):**
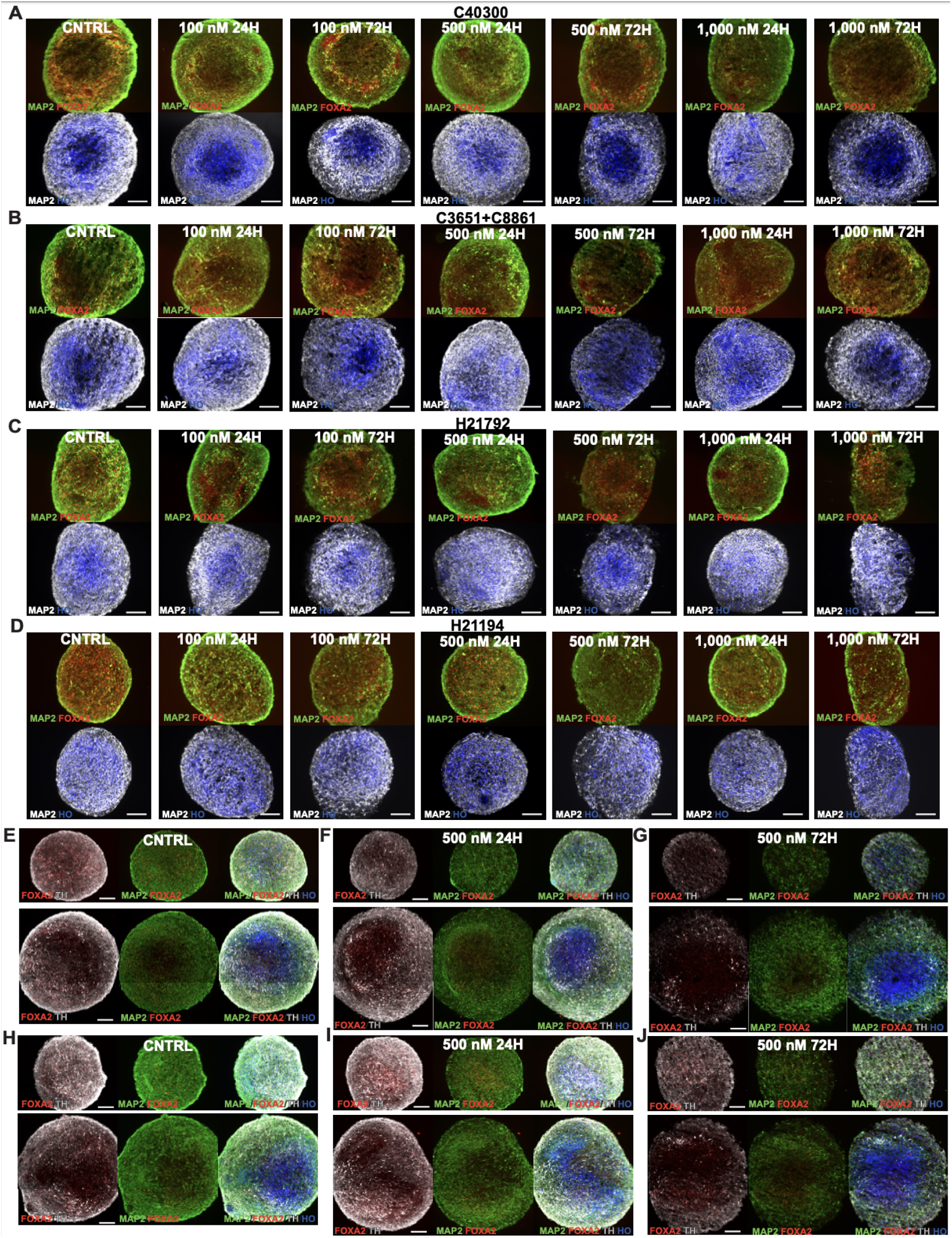
Validation and optimization of rotenone toxicity assay. A-D. Optimization of rotenone assay in chimpanzee (A-B) and human (C-D) ventral midbrain organoids, with assessment of different rotenone concentrations and durations. Immunohistochemistry for TH/FOXA2/MAP2 was used to evaluate the loss of DA neurons. E-G. Immunohistochemistry for FOXA2/TH/MAP2 in interspecies organoids (CHIR 0.6 mM) that were cultured in parallel to the organoids that were used for single cell multiome sequencing as control (E) or after 24 hours (F) and 72 hours (G) of rotenone treatment. H-J. Immunohistochemistry for FOXA2/TH/MAP2 in interspecies organoids (CHIR 0.8mM) that were cultured in parallel to the organoids that were used for single cell multiome sequencing as control (H) or after 24 hours (I) and 72 hours (J) of rotenone treatment. Scale bars: 200 μm

**Figure S9 (related to Figure 7):**
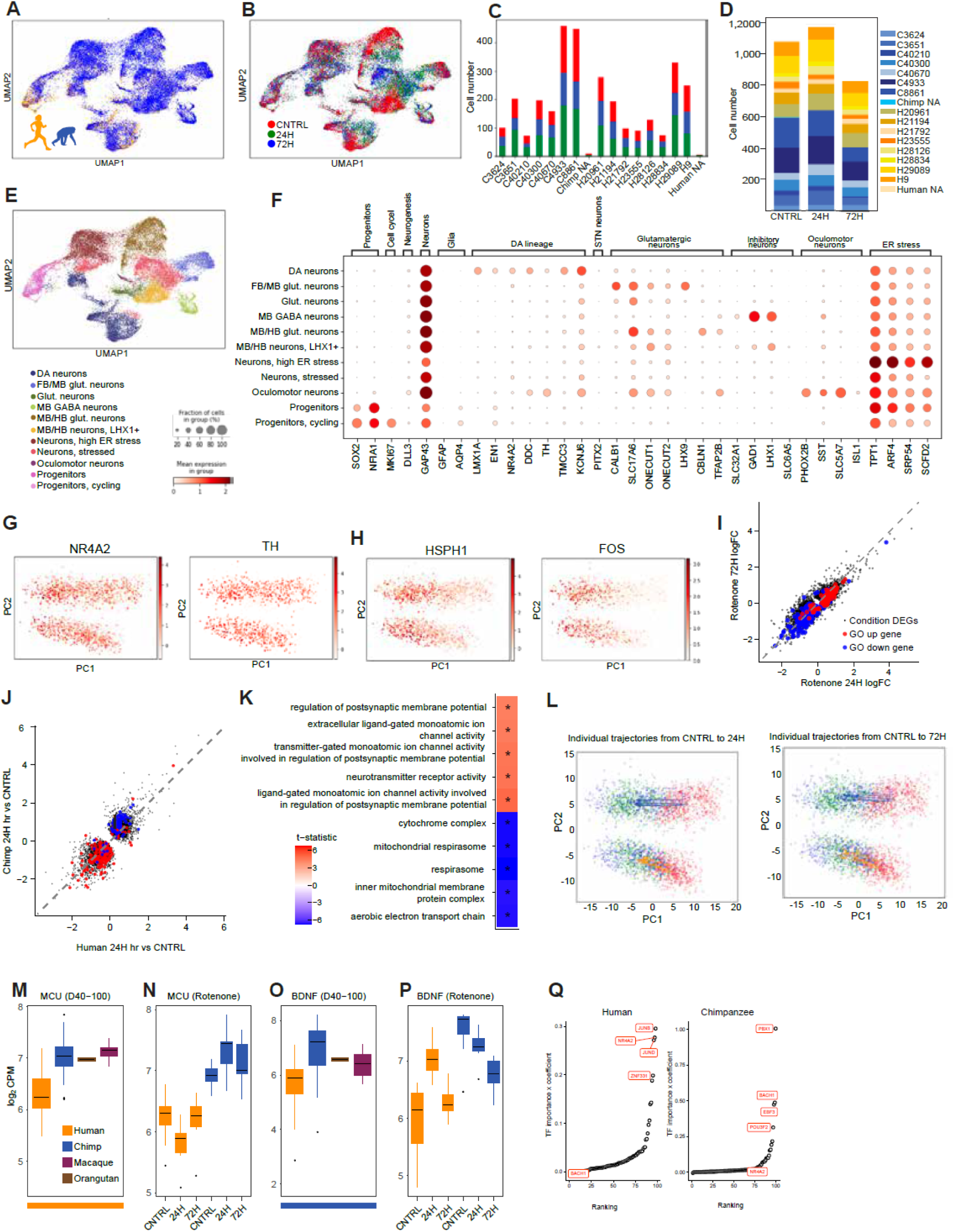
Clustering and subsetting of DA neurons in the rotenone data set. A-B. UMAPs colored by species (A) and condition (B). C. Barplot showing the number of cells per condition (control, 24 hour rotenone, and 72 hour rotenone) for each human and chimpanzee individual in the interspecies organoids that were sequenced in the rotenone experiment. D. Barplot showing the contribution of each human and chimpanzee individual in control organoids, and after 24 hours and 72 hours of rotenone treatment. E. UMAPs colored by the assigned cell type, before subsetting and further refining the DA neurons annotation. F. Dot plot of marker gene expression in the different cell type clusters G-H. PCA plot of subsetted DA neurons colored by scaled and normalized expressions of DA neuron markers *NR4A2* and *TH* (G), and heatshock protein gene *HSPH1* and early response gene *FOS* (H). I. Scatterplot showing log_2_FC for condition DEGs for 24 hour rotenone versus control and 72 hour rotenone versus control with genes in the top GO categories from Figure 7F colored and dotted y = x line. J. Scatterplot showing log_2_FC for condition DEGs for human versus chimpanzee (24 hour rotenone versus control) with genes in the top GO categories from Figure 7F colored and dotted y = x line. K. Heatmap shows the top five positive and negative gene ontology enrichment categories for species-specific rotenone (24 hours versus control) response(ranked by FDR). Positive t-statistic (red) describe categories that show a smaller magnitude condition-dependent downregulation in humans, while negative t-statistic (blue) shows categories that show a larger magnitude condition-dependent upregulation in chimpanzee. *, p < 0.05 L. PCA plots of individual cells from matched control and rotenone-exposed interspecies organoids highlight parallel trajectories related to oxidative stress in human and chimpanzee. Each vector corresponds to the distance from the centroid in the control condition to the centroid at 24 hours (left) and 72 hours (right) for each individual human (gold vector) and chimpanzee (blue vector) M. Boxplot showing normalized median *MCU* gene expression values across individuals within each species in the D40-100 dataset with colored lines below indicating polarization category. N. Boxplot showing normalized median *MCU* gene expression values across individuals within each species in the rotenone dataset at control, 24 hour rotenone, and 72 hour rotenone timepoints. O-P. Same as M-N but for *BDNF*. Q. Ranking of TF regulatory importance with directionality for BDNF in human (left) and chimpanzee (right). Top 4 TFs with the highest score of TF importance x coefficient calculated by SCENIC+ in either species are labeled.

## Supplemental information

Document S1. Figures S1–S9.

Table S1. MRI measurements of human and macaque brain volumes, related to Figure 1.

Table S2. Batch information for differentiation and sequencing, related to Figure 1 and 2.

Table S3: Differential gene expression analysis, related to Figure 4

Table S4: Differential accessibility analysis, related to Figure 5

Table S5: eGRN information in D16, D40-100, and Rotenone dataset, related to Figure 6 and 7

Table S6: Condition- and species-dependent response to rotenone, related to Figure 7

## STAR Methods

### EXPERIMENTAL MODEL AND STUDY PARTICIPANT DETAILS

#### Pluripotent stem cell lines

Human, chimpanzee, orangutan, and macaque pluripotent stem cell lines^104–109^ were maintained on recombinant human laminin-521 (0.6-1.2 µg/cm^2^, BioLamina) in StemMACS™ iPS-Brew XF medium (Miltenyi Biotec). The cells were passaged with EDTA (0.5 mM) every 3-4 days at a ratio of 1:6-1:12 for human, chimpanzee and orangutan lines and 1:15-1:30 for the macaque lines. When passaged, the pluripotent cells were seeded into StemMACS™ iPS-Brew XF medium supplemented with 3 μM Thiazovivin for 2-5 hours to improve the recovery. The cells were tested and confirmed to be negative for mycoplasma.

#### Primary midbrain samples

Macaque midbrain tissue was obtained for the developmental time points PCD40, PCD65 (n=2), PCD80 (n=2) and PCD90 from the Primate Center at the University of California, Davis. Cortical samples from the same fetal brains have previously been analyzed in Schmitz et al.^110^. All animal protocols were approved by the Institutional Animal Care and Use Committee at the University of California, Davis and all procedures were performed in accordance with the requirements of the Animal Welfare Act. Additional macaque data from PCD110 (*n* = 2) were taken from Zhu et al.^111^. One sample of de-identified human tissue was collected at GW23 with previous patient consent in strict observance of legal and institutional ethical regulations in accordance with the Declaration of Helsinki. Protocols were approved by the Human Gamete, Embryo, and Stem Cell Research Committee and the Committee on Human Research (institutional review board) at the University of California, San Francisco. Additional first trimester human first midbrain data were taken from Eze et al.^43^ and Braun et al.^42^

### METHOD DETAILS

#### Nutlin-3a sensitivity assay to determine p53 status

To exclude any cell lines with aberrant p53 expression, most of the lines included (with the exception of H21194, O11045-4593 and LYON-ES1) were screened in a Nutlin-3a sensitivity assay, as described previously^112–114^. In brief, pluripotent cells were passaged with EDTA (0.5 mM) at 1:3 and plated without rock inhibition in stem cell medium containing 10 µM Nutlin-3a (Selleck Chemicals). In pluripotent lines with a normal p53 status, this resulted in cells undergoing apoptosis within 24 hours.

#### Generation of pooled cultures of ventral midbrain progenitors

Prior to starting the differentiation, all pluripotent stem cell lines were cultured separately. To initiate the differentiation, the cells were dissociated with EDTA and counted in order to add an equal number of cells from each line to the pooled cultures. For ventral midbrain induction and cryopreservation of the ventral midbrain progenitors, the protocol described in Nolbrant el al., 2017^31^ was used. In brief, pooled pluripotent stem cells were seeded at 12,500 cells/cm^2^ (312,500 cells/T25 flask) onto recombinant human laminin-111 in N2 medium with dual SMAD inhibition (10 μM SB431542 + 100 ng/ml Noggin) from day 0-9. To obtain the correct ventral and rostro-caudal identity, 300 ng/ml SHH-C24II and 0.4-0.8 μM CHIR009921 were added between day 0-9 (when culturing the macaque lines individually a CHIR009921 concentration of 0.2-0.4 μM was used). To finetune the patterning of the ventral midbrain progenitors, 100 ng/ml FGF8b was added to the medium from day 9. On day 11, the early midbrain progenitors were passaged and replated at high density (800,000 cells/cm^2^). Between day 11-16, the cells were maintained in B27 medium supplemented with 20 ng/ml BDNF, 200 μM Ascorbic acid (AA) and 100 ng/ml FGF8b. The identity of the progenitors was determined by immunocytochemistry on day 14, before the cells were cryopreserved on day 16, or directly assembled into ventral midbrain organoids. For a complete list of differentiation experiments in this study, refer to Table S2.

#### Generation of ventral midbrain organoids

To generate ventral midbrain organoids, 10,000 day 16 progenitors were seeded into each well of a U-bottom ultra-low attachment 96-well plate. The organoids were cultured in organoid maturation media containing Neurobasal supplemented with 1x B27, 1x GlutaMAX, 1x MEM NEAA, 20 U/mL Pen-Strep, 55 μM 2-Mercaptoethanol, 20 ng/ml BDNF, 200 μM AA, 10 ng/ml GDNF and 500 μM db-cAMP^115^. For the initial seeding, 10 μM Y-27632 was also added. Two-thirds of the media was changed every 2-3 days and the organoids were moved to ultra-low attachment 6-well plates on day 40, with 10 organoids per well. During the first round of long term maturation it became apparent that one human line (H28834) kept proliferating over an extended time and that it at day 100 predominantly gave rise to a dorsal midbrain-hindbrain progenitor cell that was characterized by the expression of *PAX3/7* and *ZIC1*. To optimize the maturation and limit the expansion of these dorsal progenitors, we used organoid maturation media supplemented with 10 μM DAPT from day 40 in subsequent maturation experiments (Rotenone challenge experiment, Outgroup experiment, see Table S2).

There are several keys to our success of making multi-individual interspecies organoids. First, we screened the cells for p53 function to remove lines that would have an obvious growth and survival advantage and that might bias the results from the rotenone vulnerability assay and lead to false conclusions. Second, by using a highly efficient protocol for neuronal induction in 2D we push the cells to differentiate into midbrain progenitors. This allows us to mix up to 17 individuals at the pluripotency stage and generate pools of neuronal progenitors that can be assembled into 3D spheres for neuronal maturation. Third, at day 40, we introduce a new method to overcome imbalances, optimizing the media composition by testing different combinations of small molecules to promote cell cycle exit and neuronal differentiation.

#### Immunochemistry

To process organoids for histology, the organoids were washed in PBS and then fixed in 4% paraformaldehyde (PFA) solution for 30 min at room temperature with occasional inversion of the tube to ensure full submersion in PFA. The organoids were then washed 2x in PBS and left in 30% sucrose in PBS overnight for cryoprotection. The following day, the organoids were transferred to embedding solution containing equal parts OCT and 30% sucrose in PBS and placed in cryosectioning molds on dry ice to freeze. The organoid blocks were then stored at -80°C degrees until sectioned on a Leica Cryostat Model CM3050 S, at a thickness of 20 μm in series of 1:12 to 1:15 and mounted onto Superfrost® plus adhesion microscope slides. A minimum of 5 organoids were processed and stained for each condition. To prepare for immunohistochemistry, the slides were air-dried and then washed 1x in PBS before being placed horizontally in a humidity chamber and covered with blocking solution of either PBS + 0.2 % triton-x + 5 % donkey serum or PBS + 0.2% triton-x + 1% BSA for 1 hour. The blocking solution was subsequently removed and replaced with blocking solution containing the 1° antibodies (Rabbit Anti-Tyrosine Hydroxylase Antibody, 1:1,000; Goat Anti-Human Hnf-3 beta/foxa2 Polyclonal Antibody, 1:500; Mouse Anti-MAP2 (2a+2b) Antibody, 1:500; Mouse Anti-Human GFAP, 1:1,000; Rabbit Anti-NFIA, 1:1,000) for incubation in room temperature overnight. The following day, the slides were washed 2x in PBS and then put back in the humidity chamber for incubation in blocking solution for 30 min. After the second blocking step, the slides were incubated in blocking solution containing the 2° antibodies and Hoechst (1:10,000, ThermoFisher, 62249) for 1 hour at room temperature and then washed 3x in PBS before being cover slipped using Aqua-Mount mounting medium.

For immunocytochemistry of D16 progenitors, the cells were fixed in 4% PFA for 15 min at room temperature and then washed 2x in PBS. To block unspecific staining, the cells were incubated in either PBS + 0.1 % triton-x + 5 % donkey serum or PBS + 0.1% triton-x + 1% BSA for 1-3 hours. After removing the blocking solution 1° antibodies (Mouse Anti-HNF-3β Antibody (RY-7), 1:500; Goat Anti-Human Hnf-3 beta/foxa2 Polyclonal Antibody, 1:500; Rabbit Anti-LMX1 Polyclonal Antibody, 1:1,000; Goat Anti-Human Otx2 Polyclonal Antibody, 1:1,000) in blocking solution was added to the cells and the plates were stored at 4°C overnight. The following day the cells were washed 3x in PBS before blocking solution containing 2° antibodies and Hoechst was added. After 2 hours of incubation, the cells were washed 3x in PBS.

#### RNAscope RNA in situ hybridization

RNAscope Multiplex Fluorescent Reagent Kit v2 assay (ACD, 323120) was performed on fixed organoid 10 μm sections in accordance with the user manual provided by Advanced Cell Diagnostics (ACD, 232100-USM/Rev Date: 02272019). To detect in situ hybridization (ISH) signals, Opal^TM^ fluorophores were applied and the sections were coverslipped with ProLong™ Gold Antifade Mountant with DAPI for counterstaining of nuclei.

#### Imaging

Brightfield images of midbrain progenitor cultures and organoids were captured using an Olympus brightfield scope Model CKX53 microscope and with the image acquisition software Application Teledyne Lumenera Infinity Analyze, Version 7.0.3.1111. Fluorescent immunochemistry images were captured using an Invitrogen Evos AMF7000 microscope, using the image acquisition software Application M7000, version 2.2.804.158. Organoids used for RNAscope were imaged at high resolution on a laser scanning confocal microscope (Stellaris, Leica Microsystems) with a 40X oil (1.3NA) Plan Apochromat objective and a pixel size of 446nm x 446nm. Images from all samples were acquired under the same imaging settings and stitched automatically using the LASX software. Images were processed using ImageJ^116^ version 2.1.0/1.53c. Any adjustments made to the images were applied equally across the image, and without the loss of information.

#### Rotenone stress challenge of interspecies organoids

Before inducing oxidative stress in the interspecies organoids, we tested the sensitivity of single-species human and chimpanzee midbrain organoids to rotenone treatment. To do this, concentrations of 100 nM, 500 nM and 1,000 nM of freshly prepared rotenone were applied to the organoid maturation media for 24-72 hours, to organoids from multiple individuals of each species (after 80-108 days in culture). Based on visual assessment of TH^+^ fiber density in these organoids, a concentration of 500 nM rotenone was subsequently used for ventral midbrain interspecies organoids for the 10x multiome snRNA- and ATACseq experiment. At day 80, a full media change was performed, and the media was replaced with either organoid maturation media (control) or organoid maturation media with rotenone (stress challenge). After 24 hours, organoids for the first rotenone challenge time point were collected, and after 72 hours the control organoids and the organoids for the second rotenone challenge timepoint were collected and processed for immunohistochemistry and sequencing.

#### High resolution diffusion MRI of postmortem macaque brains

Diffusion MRI acquisition details on macaque brains can be found in a previous publication^117^. Brain samples were previously collected as part of this atlas and analyzed as follows. Three young adult postmortem macaque brains were perfusion-fixed in 4% paraformaldehyde solution in 0.1M PBS. They were immersed in 0.1M PBS for at least 120 hours before imaging and transferred to Fomblin (Fomblin Profludropolyether; Ausimont, Thorofare, NJ) just before imaging. Diffusion MRI were acquired on a Bruker BioSpin 4.7 T MRI (Bruker) machine for each sample using a 3D multiple spin echo diffusion tensor sequence (b=1000s/mm^2^; 30 directions) with the following parameters: repetition time=0.7s; echo time=32.5ms. The resolution was 0.39×0.52×0.54mm^3^. Overall scanning time for each sample was 45h.

#### T1w and diffusion MRI of in-vivo human brains

T1w and diffusion MRI data of 10 in-vivo human brains were downloaded from the human connectome project young adult (HCP-YA) dataset^118^. The imaging resolution for T1w MRI was 0.7mm isotropic. The imaging resolution for dMRI was 1.25mm isotropic.

#### Image processing, tractography and calculations of ratios

MRI data from macaque and human subjects had been collected previously, as described in above method sections, as part of the WU-Minn Human Connectome Project ^118^ and as a macaque brain atlas (co-authors of this study^117^). In the current study, these data were used to define new regions of interest (ROI) by an experienced neuroanatomist (FW) and to perform new measurements based on our macaque^117^ and human^119^ atlases. Volumes of ROIs were quantified using the number of voxels in each ROI multiplied by the product of resolution.

Whole-brain deterministic tractography^117,120,121^ was performed with Diffusion Toolkit (Version 0.6.4.1) software with a fractional anisotropy threshold of 0.2 and an angular threshold of 42 degrees. Fibers that pass through each pair of ROIs were delineated.

For each ROI, we computed the ratio between mean volumes of human and macaque and performed unequal variances Satterthwaite T-test for the ratio of two means (Rothmann, Wiens, and Chan, 2011), with the null hypothesis that the ratio of the group means is 1. The standard deviations of the ratios plotted in the figure were calculated according to a Taylor series expansion approximation (Stuart and Ord et al., 2010) using the following formula:

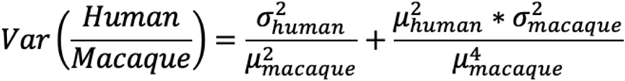

Where σ_human and σ_macaque are standard deviations of the human and macaque ROI volumes, and μ_human and μ_macaque are the means of the human and macaque ROI volumes.

Since these human/macaque ratios are already derived test statistics between two groups with unequal sample sizes, it is inappropriate to perform further statistical tests on them (such as comparing ratios across ROIs). However, from the values of the human/macaque ratio and the small standard deviation of the ratios, we can see that the ratios between ROIs are significantly different.

#### Dopamine release assay

Human and chimpanzee intraspecies pooled midbrain organoids (CHIR 0.8µM) were maintained in organoid maturation media in 48-well plates with three organoids per well. For dopamine release assessment, the media was removed, and the organoids were washed five times with 600 µL of HBSS. They were then incubated in 155 µL of HBSS at 37°C for one hour. Following the incubation, 150µL of the supernatant was harvested into Eppendorf tubes, flash frozen on dry ice, and stored at -80°C. For KCl induced depolarization, the organoids were incubated at 37°C for 30 minutes in 50mM KCl in culture media before washes and incubation with HBSS.

DA sniffers cells are a HEK-293 Flp-In T-Rex cell line stably expressing fluorescent G protein-coupled receptor-based DA sensor GRAB-DA2M^122,123^. They were maintained in DMEM supplemented with 10% FBS, 15 µg/mL blasticidin, and 200 µg/mL Hygromycin B. Cells were passaged at least twice before they were used in experiments. 24 hours before the measurements were taken, cells were seeded in imaging chambers (18-well, ibidi) coated with poly-L-ornithine in media complemented with 1 µg/mL tetracycline to induce sensor expression. Live DA sniffer cell imaging was performed on a widefield Leica microscope using a 20x NA 1.4 objective. Three images were taken per well: baseline fluorescence (F0), image after the addition of sample collected from brain organoids (Fsample), and a maximum fluorescence response after the addition of 10 µM DA solution (Fmax). Mean fluorescence intensity was extracted for each image using ImageJ (NIH) and the response was reported as %max = (Fsample-F0)/(Fmax-F0). Data was visualized using GraphPad Prism.

#### Organoid culture for multielectrode array (MEA) recordings

Human and chimpanzee organoids (CHIR 0.8µM) were transferred to recording media on day 60 to optimize physiological relevance. Recording media consisted of BrainPhys (StemCell, #05790), 1% B27 (ThermoFisher, #17504044), and 1% Amphotericin B (Gibco, #15290-026). Note: Organoids left in Neurobasal A as a control exhibited minimal activity, supporting the use of BrainPhys as a more effective medium for recordings. Additionally, organoids typically required several days in BrainPhys before showing robust activity. Immediately prior to recording, organoids were transferred via a cut-tip pipette to a MaxOne PSM chip (MaxWell Biosystems, #MX1-S-CHP) for acute recordings in minimal media. Minimal media was used to ensure optimal electrode contact in an acute setting. Sealing lids (MaxWell Biosystems, #MISC-LID-FEP) with gas-permeable, water-impermeable membranes (MaxWell Biosystems, #MX1-LID-MEM) were employed to minimize evaporation and resulting osmolarity changes. Additional media was added within a maximum of 40 minutes (typically under 25 minutes) to support long-term organoid health. If the organoid adhered to the electrode array during this period, it was subsequently cultured on the chip for chronic measurements; otherwise, it was returned to a 24-well plate for acute measurements. Data presented in Figure S3 represent chronic measurements of the same organoid across multiple timepoints.

#### MEA recordings

Recordings were conducted using the MaxOne single-well high-density MEA system (#MX1-SYS) placed inside an incubator. Activity scans were performed on the area of the MEA covered by the organoid (1,020 electrodes per configuration, scanned for 30 seconds per configuration). Recording configurations were generated based on activity scan results, with the following parameters: Method: Neural Units, Selection Preference: Firing Rate, Number of electrodes per group: 9, Maximum Number of Units: 150, Minimum Spacing Between Units: 50 µm, and Number of Electrodes to Select: 1,020. Measurements were made in a culture incubator (5% CO2 at 37°C) with a sampling rate of 20 kHz for all recordings and saved in HDF5 file format. Recordings typically lasted 5-10 minutes, depending on the amount of activity seen. If no credible activity was seen in an activity scan, the recording was skipped.

#### Spike sorting and curation

Data were preprocessed as in Sharf et al.^124^. Briefly, the raw extracellular recordings were band-pass filtered from 300-6000 Hz, then spike-sorted using the Kilosort2 algorithm through a custom Python pipeline. The resulting clusters were manually curated to remove noise and multi-unit activity. Curation was based on confirming neuronal spike waveforms, inter-spike intervals consistent with the absolute refractory period, and Gaussian amplitude distributions. In cases of duplicate clusters (defined as clusters with locations <50 µm apart and >95% of spikes concurrent), the cluster with smaller amplitude and fewer spikes was removed. Neural clusters are referred to as ‘units’ in the results section.

### Generation of single cell multiome GEX and ATAC data from pluripotent stem cell-derived neuronal progenitors and organoids

#### Preparation of samples for sequencing

Two types of samples were used for 10x Genomics single nuclei multiome GEX + ATAC sequencing: flash frozen D16 progenitors and ventral midbrain organoids (D40, D80 and D100). To prepare the D16 progenitors, the cells were dissociated for 10 min in Accutase, counted and distributed into Eppendorf tubes at 1-2 million cells per tube. The cells were centrifuged at 400 x g for 5 min and all liquid was removed before the tubes were dipped into an isopentane bath on dry ice for flash freezing. To prepare the organoid samples, organoids were cut into smaller pieces under a dissection microscope using scalpels (halves or quarters depending on timepoint) and then transferred to Eppendorf tubes. For the D40 time point, 20 organoids were placed in each tube and for D80-100, 10-15 organoids were used. The tubes were centrifuged at 100x g for 2 min to remove all the liquid and then dipped into an isopentane bath on dry ice for flash freezing.

#### Nuclei isolation for mutiome sequencing

To isolate the nuclei from D16 progenitors, the 10x Genomics demonstrated protocol *Nuclei Isolation for Single Cell Multiome ATAC + Gene Expression Sequencing (CG000365 - Rev B)* was used with a few modifications. The cells were incubated for 4 min on ice in 0.1x lysis buffer and washed in wash buffer containing 2% BSA. To isolate nuclei from organoids, the 10x demonstrated protocol *Nuclei Isolation from Embryonic Mouse Brain for Single Cell Multiome ATAC + Gene Expression Sequencing (CG000366 - Rev C)* was used and the organoid pieces were incubated for 5 + 5 min on ice in 0.1x lysis buffer and then washed in wash buffer containing 2% BSA. For the D80 and D100 organoid samples, two additional washes were added to the procedure to remove as much debris as possible. After the final wash, nuclei from both D16 progenitors and organoids were passed through a 40 μm Scienceware® Flowmi™ cell strainers and the number of nuclei and the viability was determined by Propidium iodide (PI)/HOECHST or Trypan blue staining using a Invitrogen Countess Model Countess 3 FL. The quality of the nuclei was further verified using a microscope with a high magnification 40x objective in order to assess the integrity of the nuclear membranes and the amount of debris present, as recommended by 10x Genomics.

#### Library generation and sequencing

To process nuclei and to generate multiome single cell GEX and ATAC libraries, the 10x Genomics provided protocol *(Chromium Next GEM Single Cell Multiome ATAC + Gene Expression User Guide CG000338 Rev C)* was used. We loaded Chromium NEXT GEM Chip J with a target for recovering 10,000-21,000 nuclei per lane and ran on the Chromium controller. The quality of the resulting libraries was assessed using the 2100 Bioanalyzer and high sensitivity DNA kits (Agilent) before they were sequenced on the Illumina NovaSeq S4 platform with the recommended cycle parameters. For details on 10x collection batches and sequencing metrics, see Table S2.

#### MULTIseq labeling, library preparation and sequencing

To allow for multiplexing of samples during 10x sequencing and subsequent demultiplexing of sample identity, MULTIseq labeling^36^ was used during the two first sequencing runs. The MULTIseq workflow was nearly identical to what has been described by McGinnis et al.^36^ with some minor modifications. The labeling was performed based on CHIR concentration after the nuclei had been isolated and quantified. Nuclei from different CHIR conditions were then mixed and transposed according to the 10x Genomics protocol. During labeling, the nuclei were incubated with the barcode-Anchor LMO mix at a concentration of 50 nM and subsequently incubated with the Co-Anchor LMO at 50 nM. To prepare the MULTIseq library, 10 μl of the product from ‘Pre-Amplification SPRI Cleanup’ (Step 4.3) portion of the 10x library construction protocol was used. The MULTIseq libraries were sequenced using the Illumina NovaSeq S4 platform and sample classifications were performed using the method described by https://github.com/mtvector/py-multi-seq.

### Analysis of single-cell multiome datasets

#### Reference genomes and read alignment

Our reference data were aligned and annotated in a prior study^45^. The authors aligned many human and nonhuman primate genomes using Progressive Cactus^125^ and annotated them using the Comparative Annotation Toolkit (CAT)^126^ with the human Gencode annotation v33^127^ as the source for homology-based annotations. The genome assembly versions that we used from this alignment were hg38 (human), panTro6 (chimpanzee), ponAbe3 (orangutan), and rheMac10 (rhesus macaque). As part of its alignment strategy, Progressive Cactus imputes the genome of the most recent common ancestor of each group of genomes it aligns, and we chose to use the *Homo/Pan* most recent common ancestor (MRCA) for snRNA-seq alignment to mitigate reference bias toward either the human or chimpanzee lineage.

We have found that some CAT annotations mistakenly place certain mitochondrial genes on nuclear contigs, possibly due to nuclear mitochondrial insertions (NUMTs). We also observed that the imputed *Homo/Pan* MRCA genome lacked a complete mitochondrial sequence. To remedy this, we constructed a consensus mitochondrial sequence by aligning the human, chimpanzee, bonobo, and orangutan mitochondrial genomes using MUSCLE^128^ and choosing the most common base at every position, using orangutan to break ties. We then removed from the *Homo/Pan* MRCA annotation all genes whose human source transcript is mitochondrial in the Gencode human annotation, and then lifted all of the Gencode mitochondrial annotations to the consensus mitochondrial sequence using liftOff^129^. Finally, we removed from the annotation genes and transcripts with the tag field set to readthrough_gene, readthrough_transcript, PAR, fragmented_locus, or low_sequence_quality. Code used to filter the gene annotation is available online: https://github.com/nkschaefer/litterbox.

We also hard-masked NUMTs in all reference genomes, using the previously described strategy^130^ of simulating reads from the mitochondrial genome, aligning to a version of the reference genome lacking the mitochondrial sequence, and calling peaks with MACS2. We then set all bases within these peaks in each reference genome to N using bedtools maskfasta^131^. This avoided calling erroneous ATAC peaks in the nuclear genome due to mismapped mitochondrial reads, and it also improved our ability to perform quality control in RNA-seq by more accurately quantifying mitochondrial gene expression.

We sought to create a map of genomic regions problematic for ATAC-seq data in chimpanzee, macaque, and orangutan, but without the use of ChIP-seq input data as has been used to produce a similar blacklist for the human genome^132^. To this end, we ran GenMap^133^ to obtain blacklists of regions where runs of at least 1kb were not uniquely mappable using 50-mers. Code to generate similar BED files of NUMT locations and unmappable regions is available online: https://github.com/nkschaefer/numty-dumpty.

#### Demultiplexing of species identity

Cells were assigned to a species of origin after counting occurrences of k-mers unique to each species’ transcriptome per cell in transcriptomic data. We downloaded each species’ CAT annotation from a prior study^45^ and then converted each annotation to FASTA format using gffread^134^. We then used FASTK [https://github.com/thegenemyers/FASTK] to count 27-mers in each transcriptome and extracted a list of k-mers unique to each species from the output. After counting the number of k-mers specific to each species’ transcriptome per cell in our data set, we were able to assign cells to species of origin by fitting a mixture model. An updated version of the method we used to assign species to cells is available online at https://github.com/nkschaefer/cellbouncer.

#### Generation of whole genome sequencing data

To generate whole genome sequencing data from human iPSC cell lines (H28126, H23555, H29089, H21792, H28834, H21194, and H20961), we purified genomic DNA from 3-5 million frozen cells (New England Biolabs, cat. T3010S)] and generated libraries using a PCR-free kit (New England Biolabs, cat. E7435L) according to the provided protocol for high sample inputs with 450 bp insert size. Libraries were sequenced on the Illumina NovaSeq S4 platform with the following cycle parameters: 151×12×24×151.

#### Demultiplexing of cell line identity

To demultiplex human cell lines, we combined the whole-genome sequencing data that we generated with previously published data for H9^135^. We aligned all data to the reference using minimap2^136^ with arguments -ax -sr, and then called variants using freebayes [<u>arXiv:1207.3907</u>] with arguments -w -0. We filtered the resulting VCF to include only biallelic SNPs with no missing genotypes using the command bcftools view -m 2 -M 2 -v snps -i ‘F_MISSING == 0’^137^. We then demultiplexed by genotype using CellBouncer demux_vcf [https://github.com/nkschaefer/cellbouncer] with a prior doublet probability of 0.2.

To demultiplex chimpanzee cell lines, we obtained bulk RNA-seq data from cardiomyocytes derived from cell lines C8861, C3624, C40300, C4933, C3651, and C40210 produced in a prior study^138^ and aligned it to panTro6 using STAR^139^ with arguments –outSAMtype BAM sortedByCoordinate –clip3pAdapterSeq AGATCGGAAGAGCACACGTCTGAACTCCAGTCA. We then added read groups to mark samples using PicardTools AddOrReplaceReadGroups [http://broadinstitute.github.io/picard/] and called variants using freebayes, limiting to genic regions, with parameters -t [BED file] -0 –min-coverage 5 –limit-coverage 100 –gvcf –gvcf-dont-use-chunk true, where [BED file] was a merged file of all gene features in the chimpanzee CAT annotation. We also obtained whole genome sequencing data for cell line C40670, processed as described previously for human whole genome sequencing data. To mitigate batch effects between these two variant sets, we limited both VCFs to only biallelic SNPs, and then used only loci present in both sets. We then merged the VCFs using bcftools merge with -m none and then filtered the resulting VCF to remove genotypes with DP < 15, variant quality < 50, and sites with more than 25% missing genotypes. We then demultiplexed individuals using CellBouncer demux_vcf [https://github.com/nkschaefer/cellbouncer] with this VCF file and a prior doublet probability of 0.2.

To demultiplex rhesus macaque cell lines without preexisting sequence polymorphism data, we clustered mitochondrial haplotypes in rhesus macaque cells from the D16 outgroup experiment library using CellBouncer demux_mt on the ATAC-seq data [https://github.com/nkschaefer/cellbouncer], which yielded three clusters that we validated by eye using hierarchical clustering. We then inserted read groups into the RNA-seq BAM file for the same cells, labeling individual of origin, as learned from mitochondrial clustering, via the sample field. For this step, we only used cell-to-mitochondrial haplotype assignments with a Bayes factor (confidence metric) of at least 5. We then obtained a set of nuclear genome variants segregating across individuals by running freebayes on this BAM file with arguments -w -0, and then filtered the resulting VCF by keeping only biallelic SNPs with variant quality of at least 1000, with no missing genotypes and where all called genotypes had DP >= 15. Finally, we assigned cells to individuals of origin by running CellBouncer demux_vcf using this variant call set.

#### Pre-processing and normalization of GEX data

The single cell data analysis was performed in Scanpy, using AnnData structures^140,141^. Interspecies and interindividual doublets were removed based on the demultiplexing for each entity. In addition, cells from the human line H28834 were removed at the D100 time point, since these cells mainly gave rise to PAX3/7^+^ progenitors at this time point, and before the maturation protocol had been refined. During quality control, cells with more than 15% mitochondrial reads or more then 10% ribosomal reads were removed. Cells were also filtered based on the number of genes (< 1,000 or > 7,500 for D40-100, and <500 or > 7,500 for D16) and reads (< 1,000) and any gene detected in less than 10 cells was removed. Post quality control and filtering, the number of cells retained for each time point was: D16=73,077; D40=67,416; D80=34,107; and D100=4,254 cells. Counts in cells were then normalized by read depth (CPM normalization), log transformed and then scaled to unit variance and zero mean, with a max value of 10. Principal component analysis (PCA) was performed using the top 15,000 most variable genes and the top 50 principal components were computed.

#### Cross-species integration and clustering of GEX data

To obtain clusters with merged species identities, Batch Integration of batch-balanced k-nearest neighbors (BBKNN) was applied^142^. The size of the local neighborhood was set to find 3 neighbors per batch and the batch_key was set to species. Leiden clustering using BBKNN-derived k-nearest-neighbor graphs was then applied according to the KNN graph with the resolution parameter set to 1.5. After the initial clustering, leiden clusters in D40-100 organoids defined by high ER stress (*TPT1*, *ARF4*, *SRP54* and *SCFD2*)^38,39^, poor QC metrics, and a lack of cell type markers were removed. The same process of PCA, BBKNN and leiden clustering as outlined above was then performed again. The resulting leiden clusters were classified with supervised names and merged according to subtype identity as informed by cluster-enriched genes and known cell type and neuronal subtype markers from the literature.

#### Cross-species integration and clustering of GEX data, rotenone data set

Because rotenone exposure reduced cell type marker gene expression and made it more difficult to delineate homologous cell types across species, we used a slightly different approach to integrate and cluster the rotenone data set. We used Seurat v5.1.0^143^ and applied the same quality control parameters mentioned above as well as normalization, variable feature identification, scaling, and PCA with default Seurat parameters. Next, we performed reciprocal PCA (rPCA)-based integration to integrate data from each sequencing library for each species and rotenone time point using Seurat default rPCA parameters. We identified clusters using the default shared nearest neighbor clustering algorithm with resolution set to 1. The resulting clusters were then classified with supervised names. Finally, we subclustered the DA neuron clusters, re-integrated using rPCA with k.weight = 90, and identified clusters with resolution = 0.4. We classified cells as DA neurons if they belonged to subclusters with marker expression of established DA markers, including *TH*, *NR4A2*, *EN1* and *LMX1A*.

#### Consensus non-negative matrix factorization (cNMF) simulations

To determine the number of components, k, with the highest stability and lowest error, the data was subset to 2,000 high-variance genes and a cNMF object was generated using the cNMF command from the cnmf package with 200 iterations and a range of k values between 9 and 21. Based on the k selection plot and local density histogram, 19 components and a density threshold of 0.3 were chosen to generate the consensus object for further analysis. To identify gene expression programs (GEPs) correlated with rostrocaudal progenitor cell type identities, the normalized usages of each GEP were projected onto the UMAP for visualization and the top 100 genes contributing to each GEP were analyzed for key marker genes. To compare the distributions of normalized usage values for each GEP linked to rostrocaudal identity (Usages 3, 4, 5, 6, and 8) across CHIR concentrations, the dataset was subset to the 0.4, 0.6, and 0.8 µM CHIR conditions and the empirical cumulative distribution function (ECDF) was calculated for the normalized usage values for each GEP for each CHIR concentration using the ECDF function from the statsmodels.distributions.empirical_distribution package with default parameters. Finally, to test for significant differences in the ECDFs of GEP usage across CHIR concentrations, the two-sample Kolmogorov-Smirnov test ks_2samp function from the scipy.stats package was used with default parameters to calculate the K-S statistic and p-value for each unique pair of conditions.

#### Single-cell RNA sequencing tissue processing of primary tissue

Primary samples were prepared for sequencing as previously described by Schmitz et al.^110^. In brief, the midbrains were dissected from primary tissue samples in PBS under a stereo dissecting microscope (Olympus SZ61) and cut into smaller pieces before incubation in a prewarmed solution of papain (Worthington Biochemical Corporation). After 30–60 min of incubation in the papain solution, the samples were gently dissociated through trituration with glass pipette tips, and when needed, samples were further spun through an ovomucoid gradient to remove debris. The cells were then centrifuged at 300 x g and resuspended in PBS with 0.1% BSA and the single cell sequencing was done using the 10x Genomics Chromium controller and version 2 or 3 3-prime RNA capture kits, in accordance with 10x provided protocols. Most samples were loaded at approximately 10,000 cells per well and gene expression libraries were prepared using the 10x Genomics RNA library preparation kit and sequenced on Illumina HiSeq and NovaSeq platforms.

#### Comparison to primary midbrain data and pseudotime trajectory analysis

A public dataset of developing human midbrain described at Braun et al.^42^ was annotated and used for reference mapping and comparison with the human organoid data, as well as newly collected scRNA-seq datasets of developing human and macaque midbrain around midgestation, generated in this study. Metadata and Cellranger output aligned to hg38 were obtained from Braun et al.^42^, and cells that originated from the midbrain were subset, reclustered and annotated based on literature-supported marker expression profiles. A reference model was built with scvi-tools^144^, using the top 2,500 variable genes. Subsequently, we performed label transfer to (1) the human D40-100 organoid dataset generated in this study, and (2) the human and macaque developing midbrain datasets obtained from Eze et al.^43^ and newly generated in this study to examine correspondence of cell type assignment based on the fetal data and concordance of species specific gene expression changes in the developing midbrain, respectively. To map organoid data generated in this study to primary references and assess corresponding timepoints in midbrain development, raw counts from both primary and organoid human datasets were randomly downsampled to 50,000 cells to ensure equal representation of primary and organoid derived populations. We then performed normalization and rPCA based integration to obtain a unified UMAP for all datasets and Monocle3 to impute pseudotime trajectory.

To infer species differences in DA neuron maturation, the DA lineage (vMB progenitors, immature DA/STN neurons and DA neurons) from organoid data of all four species across three timepoints (D40, D80, D100) was subset, integrated with rPCA and Monocle3 was used to calculate pseudotime along the lineage maturation and compare between species.

#### Differential gene expression analysis in D16 and D40-100 datasets

We used dreamlet v1.3.1 for differential gene expression analysis^48,49^ with the following parameters in processAssays(): min.count=5, min.cells = 5, min.prop = 0.25, norm.method = ’RLE’. To explore major sources of variation in the data, we used variancePartition^47^ v1.35.3 with the following model formulas:

D16: ∼ (1|species) + (1|experiment) + (1|indiv) + (1|sex) + (1|pool_type) + (1|lane)

D40-100: ∼ (1|species) + (1|experiment) + day + (1|indiv) + (1|sex) + (1|pool_type) + (1|lane)

We used the following model formulas for dreamlet analysis:

D16: ∼ (1|species) + (1|experiment) + (1|indiv) + (1|sex)

D40-100: ∼ (1|species) + (1|experiment) + day + (1|indiv) + (1|sex)

where day is a continuous variable describing total days in vitro. For the alternate model described in Figure S6P, we substituted the per-cell Monocle3 pseudotime values instead of day in the D40-100 model for immature DA/STN neurons and DA neurons. Contrasts were defined as human - chimpanzee, human - macaque, and chimpanzee - macaque. Human-chimpanzee DEGs were defined as genes with FDR < 0.05 for the human versus chimpanzee contrast for each cell type. We excluded ribosomal genes from the analysis by removing all genes included in GO terms including “ribosom,” and we removed cell types with fewer than 200 human or 200 chimpanzee cells. P-values were adjusted with the Benjamini–Hochberg method across remaining cell types and genes to control FDR. To plot the correlation between DEG scores across cell types, we defined a score (logFC * -log10pval) for each gene that was DE in at least one of the plotted cell types and calculated the Pearson correlation between cell types. The score was set to NA for genes that did not meet the expression threshold in a particular cell type and the pairwise correlations between cell types were computed using only complete pairs of scores. For UpSet plots, we used ggupset v0.4.0.9 and calculated deviation as previously described^145^ with 1000 iterations. We polarized human-chimpanzee DEGs for each cell type according to the following criteria applied to adjusted p values: Human-specific: human vs chimpanzee FDR < 0.05, human vs macaque FDR < 0.05, and sign(human vs chimpanzee logFC) = sign(human vs macaque logFC); Chimpanzee-specific: human vs chimpanzee FDR < 0.05, human vs macaque FDR > 0.05, sign(human vs chimpanzee logFC) ≠ sign(human vs macaque logFC); Divergent: human vs chimpanzee FDR < 0.05, human vs macaque FDR < 0.05 and not included in human- or chimpanzee-specific; Other: not included in the other three categories. For gene set enrichment analysis, we used zenith v1.6.0, which calculates enrichment based on the full spectrum of test statistics from dreamlet analysis and we used the June 2024 version of the GO database. To calculate DA/STN lineage specificity scores (among human-specific and divergent human-chimpanzee DEGs in DA neurons and immature DA/STN neurons), we used the dreamlet function cellTypeSpecificity and summed scores for each human-chimpanzee DEG across the following D40-100 cell types: DA neurons, STN neurons, immature DA/STN neurons, Ventral FB/MB progenitors, Ventral FB MB progenitors, (cycling).

#### Differential gene expression analysis in rotenone dataset

For the rotenone dataset, our differential expression analysis approach was similar to that described above for the D16 and D40-100 datasets. Here we modeled only DA neurons and we used the following parameters in dreamlet processAssays(): min.count=5, min.cells = 5, min.prop = 0.15, norm.method = ’RLE’. We used the following model formulas for dreamlet analysis:

∼ 0 + species_condition + (1|indiv) + (1|sex)

where species_condition has values for each combination of species (human or chimpanzee) and condition (control, rotenone 24 hour, rotenone 72 hour). Contrasts were defined as:

Effect of condition in human at 24 hours: Human24H = human_24H - human_CNTRL

Effect of condition in chimpanzee at 24 hours: Chimpanzee24H = chimpanzee_24H - chimpanzee_CNTRL

Effect of condition at 24 hours: Condition24H = ((human_24H - human_CNTRL) + (chimpanzee_24H - chimpanzee_CNTRL))/2

Interaction term for species and condition at 24 hours: Species24H = (human_24H - human_CNTRL) - (chimpanzee_24H - chimpanzee_CNTRL)

Similar terms were defined for the 72 hour timepoint. Condition DEGs (contrasts beginning with “effect of condition”) were defined as genes with FDR < 0.05. Species-specific response DEGs were defined as genes with FDR < 0.1 for the contrast representing the interaction term. We used zenith for gene set enrichment analysis to examine the conserved effect of condition on gene expression at 24 hours of rotenone treatment (Condition24H contrast). To make the scatterplot of species-specific response DEGs, we took the difference of log_2_cpm values in 24 hour and control pseudobulk samples for each individual and calculated the mean across these differences for each gene in each species.

#### Peak calling, enhancer-driven gene regulatory networks (eGRNs) analysis, and in silico perturbation

The SCENIC+^64^ workflow was used to infer eGRNs in each species. First, MACS2^146^ was used for peak calling in each cell type for each species. A consensus peak set for each species was generated using the TGCA iterative peak filtering approach following the pycisTopic workflow. Each peak was extended for 250bp in both directions from the summit and then iteratively filtered to remove less significant peaks that overlapped with a more significant one. Next, the merged consensus peaks were summarized into a peak-by-nuclei matrix for each species. Topic modeling was performed on the matrix by pycisTopic using default parameters, and the optimal number of topics (15) was determined based on 4 different quality metrics provided by SCENIC+, including log likelihood. We applied three different methods in parallel to identify candidate enhancer regions by selecting regions of interest through (1) binarizing the topics using the Otsu method; (2) taking the top 3,000 regions per topic; (3) calling differentially accessible peaks on the imputed matrix using a Wilcoxon rank sum test (logFC > 1.5 and Benjamini–Hochberg adjusted P values < 0.05). To assess whether the candidate enhancers were linked to a given TF, Pycistarget and discrete element method (DEM) based motif enrichment analysis were implemented. Next, eGRNs, defined as TF-region-gene network consisting of (1) a specific TF, (2) all regions that are enriched for the TF-annotated motif, (3) all genes linked to these regions, were determined by a wrapper function provided by SCENIC+ using the default parameters. eGRNs were then filtered using following criteria: (1) Only eGRNs with more than ten target genes and positive region-to-gene relationships were retained; (2) eGRNs with an extended annotation was only kept if no direct annotation is available; 3) Only genes with top TF-to-gene importance scores (rho > 0.05) were selected as the target genes for each eGRN. Specificity scores were calculated using the RSS algorithm based on region-or gene-based eGRN enrichment scores (AUC scores). eGRNs were separately identified in human and chimpanzee, in 3 different datasets (1) D16, (2) D40-100 and (3) D80 rotenone treatment dataset. Because eGRN inference is more challenging in identifying repressive interactions^64^, we chose to focus only on eGRNs of transcriptional activators for downstream analysis. The resulting eGRN targets of transcription activators were then used to intersect with species specific DEGs and DARs and Fisher’s exact test was used to identify activator eGRNs that significantly overlapped with upregulated genes or peaks in human or chimpanzee in the DA lineage. To visualize the eGRNs identified in the D40-100 dataset, we used the method outlined in Li et al.^147^, where weighted UMAP embeddings were calculated based on TF co-expression and co-regulatory networks. Visualization of eGRN target information was done using the Pando workflow^148^ and the nodes were pruned to include only highly variable genes and peaks, DEGs or DARs.

#### Cross-species consensus peak set

We developed a publicly available tool to generate cross-species consensus peak sets, CrossPeak v1.0.0 (https://github.com/jenellewallace/CrossPeak). We were inspired by HALPER^149^, but our tool differs in that it prioritizes precise summit localization centered within fixed-width peaks and defines species-specific peaks in addition to a consensus set of orthologous peaks, among other details. Beginning with a single peak set of 501-bp summit-centered peaks for each species (human, chimpanzee, and macaque in our case but CrossPeak can handle any number of species), we used halLiftover^150^ to lift summits to the Catarrhini ancestral genome^45^. In preliminary tests, we found that shorter summit widths offered the best compromise between precise summit localization and liftover precision (success of reciprocal liftovers and minimization of liftovers that failed or went to multiple locations), so we chose 11-bp summits. Next, we took the union of the lifted ranges and excluded peaks that failed liftover, lifted to multiple chromosomes, or whose lifted summit widths were more than an indel tolerance (5 bp) different from the original summit widths. Then, we rebuilt peaks on the ancestral genome by extending the summits by 250 bp in each direction. To explore parameters for peak merging, we plotted histograms of the peak overlap widths for each pair of species and chose a value for the overlap tolerance (300 bp) that marked the transition from a nearly uniform to a nonlinearly increasing distribution. To obtain a preliminary consensus peak set, we merged human and chimpanzee peaks that overlapped by more than the overlap tolerance by taking the average summit position. Macaque peaks were added to the consensus set only if they did not overlap human-chimpanzee peaks by more than the overlap tolerance. Consensus summits were then lifted to each individual species genome. Summits were discarded if their lifted widths exceeded the indel tolerance or if they were not located close enough to the originally defined summit (within the indel tolerance for original summits or within the indel tolerance per 10 bp of summit adjustment for peaks that were merged across species). Then, peaks were rebuilt on each individual species genome by extending the summits in 250 bp in each direction, and these peaks represented round 1 consensus peaks. To capture peaks that failed to meet these strict criteria and to define species-specific peaks, we developed an additional round 2 procedure. In this case, we used halLiftover to lift longer summits (51-bp) as well as the full 501-bp peaks directly between each pair of species genomes. Peaks with summits that lifted successfully to other species within predefined indel tolerances (low = 5 bp, medium = 50 bp, high = 450 bp) or whose summits failed to lift but where the entire peak lifted successfully were added to the final consensus peak set with metadata indicating liftover accuracy. Peaks that failed to lift from the originating species to all other species’ genomes (summit and peak both failed liftover, summit lifted over but with an insertion longer than the peak width, or summit failed to lift and the peak lifted with an insertion longer than the peak width) were classified as “species-only”. Peaks that multi-mapped to any other species’ genome were excluded. Peaks that overlapped blacklisted regions, were located on non-standard chromosomes, or were located on chrM or chrY for any species were excluded for all species. For human, we used the blacklist “blacklist_hg38_unified” included in the Signac package. For chimpanzee and macaque, we used custom blacklists created as described above. Overall, CrossPeak retained 87.5% and 86.4% of the original human peaks and chimpanzee peaks, respectively.

#### Analysis of snATAC-seq data

We used Signac v1.13.9003^151^ to quantify counts for each species for the original peak set called for each species (after merging across cell types), CrossPeak consensus peaks, and CrossPeak species-only peaks in each species’ own genome coordinates. We only included cells that passed snRNA-seq quality filters and had cell type identities assigned based on snRNA-seq. We filtered cells (quality metrics calculated based on each species’ original peak set) to retain cells with TSSenrichment (as calculated with Signac function TSSEnrichment()) > 2 and counts > 1000. To categorize peaks based on their genome annotation category, we used the functions GetTSSPositions from Signac and transcriptsBy, exonsBy, and intronsByTranscript from GenomicFeatures v1.56.0. We defined promoters as extending 1000 bp upstream of the TSS. We removed duplicate annotations (with the same range and same category but different ID) from the annotation file for each species and in the case of overlaps with multiple categories, assigned each peak to the category with the greatest overlap width. To integrate data across species, we used Signac’s batch integration pipeline with CrossPeak consensus peaks (as a multi-species workaround, coordinates for each species had to be renamed to human coordinates but counts were unaffected). We used FindIntegrationAnchors() with dimensions 2 to 30 and k.anchor = 20 and IntegrateEmbeddings() with LSI reduction. We used Signac CoveragePlot() to create genome browser-style plots and the annotations are the same as those used for RNA-seq unless otherwise noted.

#### Differential accessibility analysis

We used dreamlet v1.3.1 for differential accessibility analysis of CrossPeak consensus peaks in D40-100 cell types: DA neurons, immature DA/STN neurons, and Ventral FB/MB progenitors. To improve the normalization, we downsampled the number of cells in chimpanzee and macaque pseudobulk samples using linear interpolation to match the entire range of cell numbers in human pseudobulk samples. We used the following parameters in processAssays(): min.count = 2, min.cells = 30, min.prop = 0.25, norm.method = ’RLE’. We used the following model formulas for dreamlet analysis:

DA neurons: ∼ (1|species) + (1|indiv) + (1|sex) + (1|stage) + log_num_peaks

Immature DA/STN neurons and Ventral FB/MB progenitors: ∼ (1|species) + (1|indiv) + (1|sex) + (1|day_10x) + log_num_peaks

where stage is defined as “early” for D40 and “late” for D80-100 (based on separation of D40 from D80-100 on the UMAP), day_10x represents 10X runs (collection batches) #2 and #4 from TableS2, and log_num_peaks represents log2 of the number of detected peaks per pseudobulk sample. We used “day_10x” instead of “stage” for the immature neurons and progenitors because these cell types were mostly present only at D40. CrossPeak species-only peaks could not be included in the dreamlet model since their counts were not defined in all species, but we applied the same processAssays() parameters to human or chimpanzee count matrices separately and only retained species-only peaks for analysis that met these criteria (for DA neurons).

Due to the sparsity of the ATAC data and lower power of the dreamlet model, we relaxed the significance threshold and defined human-chimpanzee DARs as peaks with adjusted p values < 0.1 for the human versus chimpanzee contrast (human-chimpanzee). P-values were adjusted with the Benjamini–Hochberg method within each cell type separately. To quantify the enrichment of evolutionary signatures overlapping DARs in DA neurons, we applied Fisher’s exact test to contingency tables of the number of overlaps between the set of human DARs and species-only peaks and evolutionary signatures defined on hg38 (except for enrichment of hDels defined on panTro6, for which we considered chimpanzee DARs and species-only peaks). Because we allowed for overlapping peaks in the CrossPeak consensus set to ensure capture of neighboring peaks that may differ across species, we first split overlapping peaks by adding half of the overlap width to each overlapping peak to create a set of nonoverlapping peaks spanning the same genomic range before calculating enrichment. We also only considered CrossPeak round 1 high confidence round 2 peaks (peaks in the low indel tolerance category) for enrichment analysis to ensure that the enrichments were not affected by peaks that were not accurately localized across species due to large insertions or deletions. For peak annotation with GREAT v4.0.4^56,57^, we used default settings to define gene regulatory domains (“Basal plus extension”). Our test set was human-chimpanzee DARs (in human coordinates) from the dreamlet analysis of CrossPeak consensus peaks and CrossPeak-defined human-only peaks, and the background set included all CrossPeak consensus peaks that met dreamlet expression thresholds plus CrossPeak-defined human-only peaks. We also set “term annotation count” to 10 to exclude GO terms with small numbers of annotated genes. To calculate the percentage of DARs overlapping promoters and their concordance with gene expression in human, we used the human NCBI Refseq annotation and defined promoters as extending 1000 bp upstream of the TSS.

#### Linking peaks to genes

We used Cicero v1.3.9^59^ to link peaks to genes. We included peaks expressed in 1% of DA lineage D40-100 cell types (DA neurons, immature DA/STN neurons, Ventral FB/MB progenitors) in at least one species, all peaks that overlapped a promoter in any species, and all peaks that met dreamlet expression thresholds in DA neurons. We ran Cicero with the same peak list for human and chimpanzee separately with default settings to obtain sets of correlated peaks for each species. With these settings, all pairs of peaks within 500 kb will appear in the output matrix with a coaccessibility score between -1 to 1. To link peaks to genes for each species across a range of coaccessiblity score thresholds, we first filtered the Cicero output matrix to keep only connections with scores greater than the threshold. Then, we linked peaks that fell anywhere within a gene body or within a promoter region 1000 bp upstream of a TSS to the corresponding gene. Next, any distal peaks linked to gene peaks were also linked to that gene. While we show that most peaks were linked to only one gene, a small minority of peaks were linked to more than one gene. We classified peaks as linked to the same gene across species if at least one gene was linked to the same peak in each species, regardless of any other linked genes. Peaks were classified as linked to different genes if the peak was not linked to any of the same genes across species. If a peak was linked to a gene in one species but not in the other (due to coaccessibility scores not exceeding the threshold), then that peak was classified as linked only in that species. We reasoned that higher signal-to-noise would allow more reliable identification of correlations between regions with higher accessibility, so we defined concordant DARs as DARs with increased accessibility linked to upregulated DEGs, considering links defined in the species where the gene was upregulated (i.e. human concordant DARs were human-upregulated DARs linked to human-upregulated DEGs based on human Cicero results and similar for chimpanzee).

#### Euclidean distance in PC space

Vectors from control to 24 or 72 hours were created by calculating the mean position of each individual at baseline, 24 hours, and 72 hours in the first two PC dimensions. These vectors were plotted per individual starting from the mean position at baseline and pointing to the mean position after treatment (either 24 or 72 hours) on top of the PCA plot of all individuals, species, and timepoints. To compare the Euclidian distance of response in PC space, the mean position per individual and condition was calculated in the first 3 PC dimensions. We calculated the norm of the vector from control to either 24 or 72 hours and used this value to make a violin plot comparing the magnitude of the vectors across species and condition. P-values were calculated from a one-way ANOVA test.

### QUANTIFICATION AND STATISTICAL ANALYSIS

Python v3.9.7 and R v4.4.0 were used for analysis and to create the plots shown in Figures 1-2 and 7 (S2, 4, snRNA-seq data in Figure S9) and Figures 4-7 (S6, S7, and S9), respectively. Statistical analyses are described in the figure legends with additional details in the corresponding Methods details sections. Values for n (number of cells, individuals, and experiments) can be found in the main text and Table S2.

