## Supplementary material for "Interspecies Organoids Reveal Human-Specific Molecular Features of Dopaminergic Neuron Development and Vulnerability": Key Resource Table

**Key resources table**

| REAGENT or RESOURCE | SOURCE | IDENTIFIER |
|---|---|---|
| **Antibodies** | | |
| Goat Anti-Human Hnf-3 beta / foxa2 Polyclonal Antibody, 1:500 | R and D Systems | Cat# AF2400; RRID:AB_2294104; Lot#ULB102108, ULB0919101 |
| Mouse Anti-HNF-3β Antibody (RY-7), 1:500 | Santa Cruz Biotechnology | Cat# sc-101060; RRID:AB_1124660; Lot# E2522 |
| Rabbit Anti-LMX-1 Polyclonal Antibody, 1:1,000 | Millipore | Cat# AB10533; RRID:AB_10805970; Lot# 3493318, 3880889 |
| Mouse Anti-MAP2 (2a+2b) Antibody, 1:500 | Sigma-Aldrich | Cat# M1406; RRID:AB_477171; Lot# 107M4856V |
| Goat Anti-Human Otx2 Polyclonal Antibody, 1:1,000 | R and D Systems | Cat# AF1979; RRID:AB_2157172; Lot# KNO0922011, KN00918121 |
| Rabbit Anti-Tyrosine Hydroxylase Antibody, 1:1,000 | Sigma-Aldrich | Cat# AB1542; RRID:AB_90755; Lot# 3574360, 3845256 |
| Mouse Anti-Human GFAP, 1:1,1000 | Takara Bio | Cat# Y40420; RRID:AB_2833249; Lot# AL90015S |
| Rabbit Anti-NFIA, 1:1,000 | Atlas Antibodies | Cat# HPA006111; RRID:AB_1854422; Lot# 000007218 |
| **Bacterial and virus strains** | | |
| **Biological samples** | | |
| **Chemicals, peptides, and recombinant proteins** | | |
| Human recombinant laminin 521 | BioLamina | Cat# LN521 |
| Human recombinant laminin 111 | BioLamina | Cat# LN111 |
| StemMACS™ iPS-Brew XF medium | Miltenyi Biotec | Cat# 130-104-368 |
| ROCK inhibitor thiazovivin | Stem Cell Tech. | Cat# 72252 |
| Nutlin-3a | Selleck Chemicals | Cat# S8059 |
| N-2 Supplement (100X) | ThermoFisher | Cat# 17502048 |
| Y-27632 | Miltenyi Biotec | Cat# 130-106-538 |
| DMEM/F-12, no glutamine | ThermoFisher | Cat# 21331020 |
| Neurobasal™ Medium | ThermoFisher | Cat# 21103049 |
| GlutaMAX™ Supplement | ThermoFisher | Cat# 35050061 |
| StemMACS SB431542 in solution | Miltenyi Biotec | Cat# 130-106-543 |
| Noggin Recombinant human | Miltenyi Biotec | Cat# 130-103-456 |

| | | | |
|---|---|---|---|
| Recombinant Human Sonic Hedgehog/Shh (C24II) | R and D Systems | Cat# 1845-SH | |
| StemMACS CHIR99021 | Miltenyi Biotec | Cat# 130-103-926 | |
| Human FGF-8b | Miltenyi Biotec | Cat# 130-095-740 | |
| B-27™ Supplement (50X), minus vitamin | ThermoFisher | Cat# 12587010 | |
| Recombinant Human BDNF Protein | R and D Systems | Cat# 248-BDB | |
| L-Ascorbic acid | Sigma-Aldrich | Cat# A4403 | |
| Dibutyryl cAMP sodium salt | Sigma-Aldrich | Cat# D0627 | |
| Recombinant Human GDNF Protein | R and D Systems | Cat# 212-GD | |
| DAPT | R and D Systems | Cat# 2634 | |
| 2-Mercaptoethanol | ThermoFisher | Cat# 21985023 | |
| MEM Non-Essential Amino Acids Solution (100X) | ThermoFisher | Cat# 11140050 | |
| Rotenone | Sigma Aldrich | Cat# R8875 | |
| BrainPhys™ Neuronal Medium | Stem Cell Tech. | Cat# 05790 | |
| MULTIseq barcoding reagents | Gartner lab; McGinnis et al.[33] | NA | |
| Protector RNase Inhibitor | Merck | Cat# 3335402001 | |
| Tetracycline | Sigma-Aldrich | Cat# T7660 | |
| Hygromycine B | ThermoFisher | Cat# 10687010 | |
| Blasticidin | ThermoFisher | Cat# R21001 | |
| Fetal Bovine Serum | ThermoFisher | Cat# 17479633 | |
| DMEM | Thermofisher | Cat# 10564011 | |
| RNAscope™ Probe- Hs-TH-C4 | ACD Bio. | Cat# 441651-C4 | |
| RNAscope™ Probe- Hs-EN1-C2 | ACD Bio. | Cat# 527741-C2 | |
| RNAscope™ Probe- Hs-KCNJ16 | ACD Bio. | Cat# 877961 | |
| **Critical commercial assays** | | | |
| RNAscope Multiplex Fluorescent Reagent Kit v2 | ACD Bio. | Cat# 323100 | |
| Chromium Next GEM Single Cell Multiome ATAC + Gene Expression Reagent Bundle | 10x Genomics | Cat# 1000283 | |
| **Datasets** | | | |
| Human fetal single cell expression data | Braun et al.[38] | https://www.science.org/doi/10.1126/science.adf1226 | |
| Human connectome project young adult (HCP-YA) dataset | Van Essen et al.[119] | https://www.sciencedirect.com/science/article/pii/S1053811913005351 | |
| **Experimental models: Cell lines** | | | |
| Human iPSCs H28126, XY, p14+12-15 | Gilad lab; Gallego Romero et al.[107] | NA | |
| Human iPSCs H23555, XY, p16+15-17 | | | Gilad lab; Gallego Rom[illegible] al.[107] |
| Human iPSCs H20961, XY, p15+12-14 | Gilad lab; Gallego Romero et al.[107] | RRID:CVCL_HA53 | |

| | | | |
|---|---|---|---|
| Human iPSCs H21792, XX, p14+12-15 | Gilad lab;<br>Gallego Romero et al.[107] | NA | |
| Human iPSCs H29089, XX, p7+12-14 | Gilad lab;<br>Gallego Romero et al.[107] | NA | |
| Human iPSCs H28834, XX, p12+10-12 | Gilad lab;<br>Gallego Romero et al.[107] | RRID:CVCL_HA54 | |
| Human iPSCs 21194, XX, p14+16-18 | Gilad lab;<br>Gallego Romero et al.[107] | NA | |
| Human ESCs WA09 (H9), XX, P37-40 | WiCell | RRID:CVCL_9773 | |
| Chimpanzee iPSCs C8861, XY, p15+14-15 | Gilad lab;<br>Gallego Romero et al.[107] | RRID:CVCL_1G34 | |
| Chimpanzee iPSCs C3624, XY, p14+13-15 | Gilad lab;<br>Gallego Romero et al.[107] | NA | |
| Chimpanzee iPSCs C40670, XY, p9+22-24 | Gilad lab;<br>Gallego Romero et al.[107] | NA | |
| Chimpanzee iPSCs C3651, XX, p11+13-15 | Gilad lab;<br>Gallego Romero et al.[107] | RRID:CVCL_1G32 | |
| Chimpanzee iPSCs C40210, XX, p14+16-18 | Gilad lab;<br>Gallego Romero et al.[107] | RRID:CVCL_1G35 | |
| Chimpanzee iPSCs C40300, XX, P14+15-17 | Gilad lab;<br>Gallego Romero et al.[107] | NA | |
| Chimpanzee iPSCs C4933, XX, p11+12-14 | Gilad lab;<br>Gallego Romero et al.[107] | NA | |
| Orangutan iPSCs 11045-4593, XX | Haussler-Salama Lab;<br>Field et al.[112] | NA | |
| ZH26-HS16, p25 | Dunbar lab; Ostrominski et al.[108] | NA | |
| Rhesus ESCs LYON-ES1, XX, p47+9-13 | Wianny et al.[111] | RRID: CVCL_XV40 | |
| ZG15-M11-10-GFP, p20 | Dunbar lab; Hong et al.[109] | NA | |
| GRAB-DA2M sniffers cells | Klein Herenbrink et al.[123] | NA | |
| **Experimental models: Organisms/strains** | | | |
| **Oligonucleotides** | | | |
| **Recombinant DNA** | | | |
| **Software and algorithms** | | | |
| Scanpy v1.8.2 | Wolf et al.[141] | https://github.com/scverse/scanpy | |
| Anndata v0.8.0 | | | Virshup et al |
| Seurat v5.1.0 | Butler et al.[144] | https://github.com/satijalab/seurat | |
| Signac v1.13.9003 | Stuart et al.[152] | https://github.com/stuart-lab/signac | |
| Dreamlet v1.3.1 | Hoffman et al.[49] | https://github.com/GGabrielHoffma/dreamdre | |

| | | |
|---|---|---|
| variancePartition v1.35.3 | Hoffman and Schadt 2016[48] | https://github.com/GabrielHoffman/variancePartition |
| CrossPeak v 1.0.0 | This paper | https://github.com/jenellewallace/CrossPeak |
| Cicero v1.3.9 | Pliner et al.[60] | https://github.com/cole-trapnell-lab/cicero-release |
| MACS2 | Zhang et al.[147] | https://github.com/macs3-project/MACS |
| ScenicPlus | Bravo González-Blas et al.[65] | https://github.com/aertslab/scenicplus |
| pycisTopic | Bravo González-Blas et al.[65] | https://github.com/aertslab/pycisTopic |
| pycistarget | Bravo González-Blas et al.[65] | https://github.com/aertslab/pycistarget |
| CellBouncer | This paper | https://github.com/nkschaefer/cellbouncer |
| Fiji | Schindelin et al.[117] | https://fiji.sc/ |
| Pando | Fleck et al.[149] | https://github.com/quadbio/Pando?tab=readme-ov-file |
| **Other** | | |
